# Metastability, fractal scaling, and synergistic information processing: what phase relationships reveal about intrinsic brain activity

**DOI:** 10.1101/2022.01.17.476583

**Authors:** Fran Hancock, Joana Cabral, Andrea I. Luppi, Fernando E. Rosas, Pedro A.M. Mediano, Ottavia Dipasquale, Federico E. Turkheimer

**Affiliations:** Department of Neuroimaging, Institute of Psychiatry, Psychology and Neuroscience, King’s College London, London, UK; Life and Health Sciences Research Institute (ICVS), School of Medicine, University of Minho, Portugal.; Division of Anaesthesia, School of Clinical Medicine, University of Cambridge; Department of Clinical Neurosciences, University of Cambridge; Leverhulme Centre for the Future of Intelligence, University of Cambridge; Alan Turing Institute, London, UK; Centre for Psychedelic Research, Department of Brain Science, Imperial College London, London SW7 2DD; Data Science Institute, Imperial College London, London SW7 2AZ; Centre for Complexity Science, Imperial College London, London SW7 2AZ; Department of Psychology, University of Cambridge, Cambridge CB2 3EB; Department of Psychology, Queen Mary University of London, London E1 4NS

**Keywords:** Functional magnetic resonance imaging, dynamic functional connectivity, complexity, metastability, fractal scaling, integrated information, LEiDA

## Abstract

Dynamic functional connectivity (dFC) in resting-state fMRI holds promise to deliver candidate biomarkers for clinical applications. However, the reliability and interpretability of dFC metrics remain contested. Despite a myriad of methodologies and resulting measures, few studies have combined metrics derived from different conceptualizations of brain functioning within the same analysis - perhaps missing an opportunity for improved interpretability. Using a complexity-science approach, we assessed the reliability and interrelationships of a battery of phase-based dFC metrics including tools originated from dynamical systems, stochastic processes, and information dynamics approaches. Our analysis revealed novel relationships between these metrics, which allowed us to build a predictive model for integrated information using metrics from dynamical systems and information theory. Furthermore, global metastability - a metric reflecting simultaneous tendencies for coupling and decoupling - was found to be the most representative and stable metric in brain parcellations that included cerebellar regions. Additionally, spatiotemporal patterns of phase-locking were found to change in a slow, non-random, continuous manner over time. Taken together, our findings show that the majority of characteristics of resting-state fMRI dynamics reflect an interrelated dynamical- and informational-complexity profile, which is unique to each acquisition. This finding challenges the interpretation of results from cross-sectional designs for brain neuromarker discovery, suggesting that individual life-trajectories may be more informative than sample means.

**Highlights:**

- Spatiotemporal patterns of phase-locking tend to be time-invariant
- Global metastability is representative and stable in a cohort of heathy young adults
- dFC characteristics are in general unique to any fMRI acquisition
- Dynamical- and informational-complexity are interrelated
- Complexity science contributes to a coherent description of brain dynamics

## 1 Introduction

There is great anticipation that functional neuroimaging may complement current clinical phenomenology in the diagnosis of disorders of brain functioning, and provide brain-based markers for patient stratification, disease progression tracking, and prediction of treatment outcomes (Zhang et al., 2021). In this context, the investigation of the brain’s functional connectivity (FC) – as revealed by resting-state functional magnetic resonance imaging (fMRI) – holds promise for enabling tools of great clinical value, with thousands of articles per year focused on elucidating properties of normal and abnormal whole-brain functionality (Zhang et al., 2021). Static FC reveals the statistical interdependence among different brain regions using blood oxygenation level dependent (BOLD) signals (Friston,1994). However, these static measures camouflage the inherent dynamic nature of brain activity which is captured with time-varying functional connectivity, or dynamic FC (dFC). Unfortunately, the fact that fMRI may be capturing something other than BOLD signals (Drew et al., 2020; Raut et al., 2021), and in the absence of a ground truth, the hurdles to use FC metrics in the clinic are high (Woo and Wager, 2015), and considerably higher for dFC due to issues of interpretation (Lurie et al., 2020) and sampling variability (Laumann et al., 2017), although the latter has been rigorously challenged (Miller et al, 2018). Moreover, the popularity of FC and dFC methods comes with a plethora of heterogeneous methodologies derived from distinct conceptualizations of brain functioning (Bijsterbosch et al., 2020).

Candidate neuromarkers should demonstrate a high degree of reliability and ideally be robust and interpretable in terms of neuroscience (Woo and Wager, 2015). Despite efforts to assess the test-retest reliability of dFC metrics, the results remain contested (Abrol et al., 2017; Bijsterbosch et al., 2017; Choe et al., 2017; Orban et al., 2020; Vaisvilaite et al., 2021; Vohryzek et al., 2020). Common approaches to address these concerns of validity include comparison of results with null models (Battaglia et al., 2020) or replication of results in alternative datasets (Varley et al., 2020). Neuroscientific interpretation of candidate neuromarkers is enhanced with convergence of evidence from multiple sources (Woo and Wager, 2015), and together with reliability, is one of the necessary conditions to introduce neuromarkers into the clinic.

With this in mind, in this paper we took a complexity-science perspective to identify a number of diverse dFC metrics for investigation (Turkheimer et al., 2021). The existence of distinct methodologies that investigate intrinsic brain activity either from a dynamical systems perspective, from considerations of the time-evolution of the dynamical system as a stochastic process, or from an information processing perspective, compels us to confront the challenging task of piecing together a coherent description of brain dynamics consistent across the underlying theories.

Two specific metrics, metastability and integrated information, derived from bottom-up and top-down analysis respectively, hold special interest for investigation. Theoretically, metastability has been described as a subtle blend of segregation and integration among brain regions that show tendencies to diverge and function independently, with tendencies to converge and function collectively (Tognoli and Kelso, 2014). Metastability has been considered a key attribute for computational models exploring mechanisms of brain dynamics and an important indicator of healthy brain functioning (Deco et al., 2017). From an alternative but complementary perspective, integrated information (operationalized as the quantity *Φ*) has been proposed as a way of quantifying the balance between integration and segregation, and possibly consciousness (Tononi, 2004). More recent metrics of integrated information, Φ*^R^*, extends this construct to reflect the degree of synergistic and transfer information processing across brain areas (Mediano et al., 2022). Therefore, we sought to investigate if these two metrics contributed converging evidence for the processes of integration and segregation that are believed to take place as part of intrinsic brain activity.

Our objective was to develop a coherent description of brain dynamics consistent across underlying theories. Therefore, rather than investigate metastability and integrated information in isolation, we assessed them in combination with metrics originating in complexity-science, as well as metrics identified theoretically or empirically as characterizing or contributing to metastability or integrated information. Whilst the methodologies used in this study have already been individually validated against null models or with surrogate data (Battaglia et al., 2020; Honari et al., 2021; Mediano et al., 2022), there is a lack of studies where these methodologies were used to compare performance in the same subjects across multiple fMRI acquisitions. Therefore, we set out to answer the following questions: are the chosen dFC metrics representative and reliable across multiple fMRI acquisitions? Are these metrics related via their ability to capture different aspects of dFC? And finally, what are the implications of these relationships?

To address these questions, we used four resting-state fMRI acquisitions recorded on two consecutive days from 99 healthy unrelated participants from the Human Connectome Project (Van Essen et al., 2013). We performed confirmatory analysis with different parcellation schemes, considering an anatomical parcellation with and without the cerebellar regions and a functional parcellation that included the cerebellar regions.

## 2 Materials and Methods

### 2.1 Data

All data used in this study was collected for the Human Connectome Project, WU-Minn Consortium (Principal Investigators: David Van Essene and Kamil Ugurbil; 1U54MH091657) with funding from the sixteen NIH Institutes and Centers supporting the NIH Blueprint for Neuroscience Research; and by the McDonell Center for Systems Neuroscience at Washington University.

### 2.2 Ethics Statement

The Washington University institutional review board approved the scanning protocol, participant recruitment procedures, and informed written consent forms, and consented to share deidentified data.

### 2.3 Participants

We used the data from the ‘500 subject’ release but restricted our analysis to the ‘100 Unrelated Subjects’ (aged 20 to 35 years old, 54 females (Glasser et al., 2013)). A list of employed subject ID numbers and associated scan times is provided in Supplementary Table S 1.

### 2.4 fMRI data acquisition and pre-processing

Each participant underwent four scans of resting-state fMRI (rs-fMRI) collected over two experimental sessions (two scans in each session) which took place on consecutive days. The datasets acquired from all participants in each of the 4 scans are referred to as ‘runs’ 1 to 4. During each scan 1200 frames were acquired using a multiband sequence at 2 millimeters (mm) isotropic resolution with a repetition time (TR) of 0.72 seconds over the span of 14 minutes 24 seconds. Participants were instructed to maintain fixation on a bright crosshair presented on a dark background in a darkened scanning room. The two scans in each session differed only in the oblique axial acquisition phase encoding. For the first 6 subjects, the rs-fMRI runs were acquired using a Right-Left (RL) phase-encoding followed by a Left-Right (LR) phase-encoding on both days. For the following 94 subjects, the order of the different phase-encoding acquisitions for the rs-fMRI runs across days was counterbalanced (RL followed by LR on Day 1; LR followed by RL on Day 2).

Data were pre-processed with the HCP’s minimal pre-processing pipeline, and denoising was performed by the ICA-FIX procedure (Glasser et al., 2013; Griffanti et al., 2014; Salimi-Khorshidi et al., 2014). A complete description of the acquisition and pre-processing details may be found at the HCP website https://www.humanconnectome.org/. One subject was excluded from the analysis as the image file was corrupted.

### 2.5 Parcellations

We parcellated the pre-processed fMRI data by averaging time-courses across all voxels for each region defined in the anatomical parcellation AAL (Tzourio-Mazoyer et al., 2002) considering all cortical and subcortical brain areas including the cerebellum, N=116 or without the cerebellum N=90. For confirmation of the contribution of the cerebellum to the reliability of the metrics, we also parcellated the fMRI data with the NEUROMARK framework (Du et al., 2020).

### 2.6 Bandpass filtering

To isolate low-frequency resting-state signal fluctuations, we bandpass filtered the parcellated fMRI time-series for 0.01-0.08 Hz, in alignment with previous studies (Glerean et al., 2012).

### 2.7 Phase relationships

We investigated two complementary forms of phase relationships, phase-locking and phase synchrony as illustrated in Figure 1.

**Figure 1.**
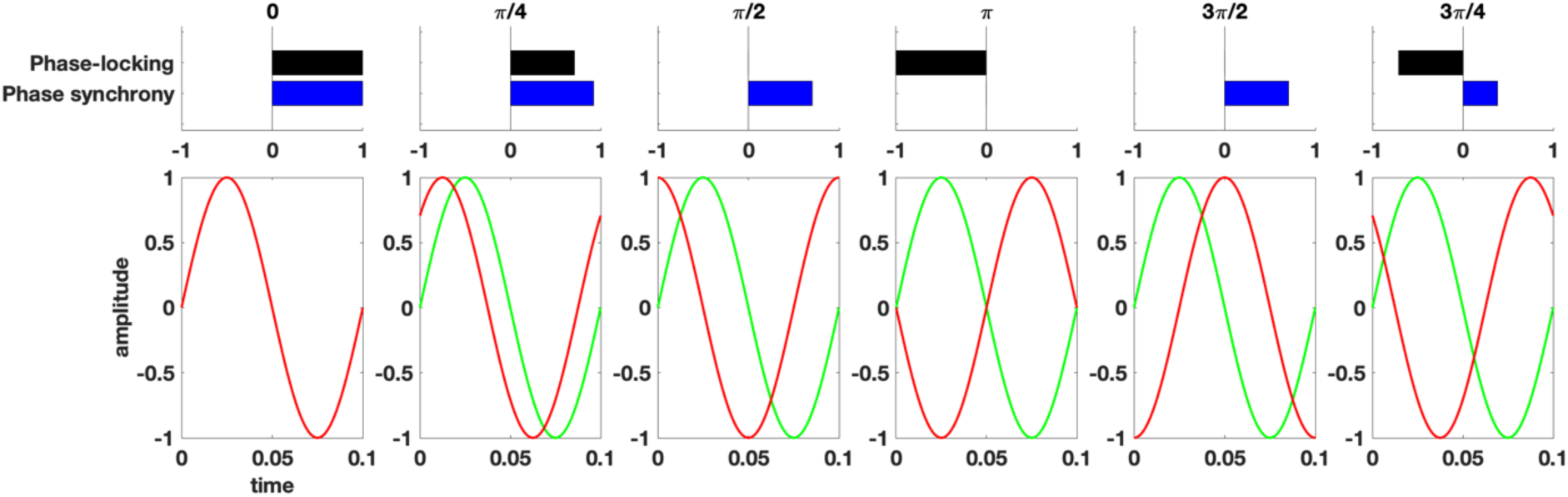
Two complementary forms of phase coupling for the calculation of dFC metrics. Phase-locking, evaluated as the cosine of the phase difference, is sensitive to both in-phase and anti-phase relationships between regions, while phase synchrony is sensitive to the phase alignment between regions.

### 2.8 Functional connectivity through phase-locking

We estimated functional connectivity (FC) with the nonlinear measure of phase-locking which may be more suitable than linear measures such as Pearson correlation for analyzing complex brain dynamics (Pereda et al., 2005; Quian Quiroga et al., 2002). Indeed, phase relationships have been leveraged in many dFC studies to date (Alonso Martínez et al., 2020; Cabral et al., 2017; Deco and Kringelbach, 2016; Figueroa et al., 2019; Ponce-Alvarez et al., 2015; Vohryzek et al., 2020; Zhang et al., 2019; Zhou et al., 2020). First, we calculated the analytical signal using the Hilbert transform of the real signal (Gabor, 1946). Then, the instantaneous phase-locking between each pair of brain regions *n* and *p* was estimated for each time-point *t* as the cosine difference of the relative phase as

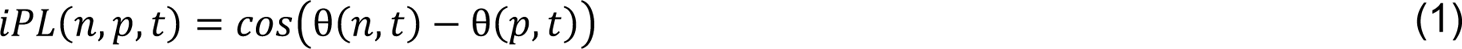

Phase-locking at a given timepoint ranges between -1 (regions in anti-phase) and 1 (regions in-phase). For each subject the resulting *iPL* was a three-dimensional tensor *NxNxT* where N is the dimension of the parcellation, and *T* is the number of timepoints in the scan.

#### 2.8.1 LEiDA – Leading Eigenvector Dynamic Analysis

To reduce the dimensionality of the phase-locking space for our dynamic analysis, we employed the Leading Eigenvector Dynamic Analysis (LEiDA) (Cabral et al., 2017) method. The leading eigenvector *V_1_(t)* of each *iPL(t)* is the eigenvector with the largest magnitude eigenvalue and reflects the dominant FC (through phase-locking) pattern at time *t*. *V_1_(t)* is a *Nx1* vector that captures the main orientation of the fMRI signal phases over all anatomical areas. Each element in *V_1_(t)* represents the projection of the fMRI phase in each region into the leading eigenvector. When all elements of *V_1_(t)* have the same sign, this means that all fMRI phases are orientated in the same direction as *V_1_(t)* indicating a global mode governing all fMRI signals. When the elements of *V_1_(t)* have both positive and negative signs, this means that the fMRI signals have different orientations behaving like opposite anti-nodes in a standing wave. This allows us to separate the brain regions into two ‘communities’ (or poles) according to their orientation or sign, where the magnitude of each element in *V_1_(t)* indicates the strength of belonging to that community (Newman, 2006). For more details and graphical representation see (Figueroa et al., 2019; Lord et al., 2019; Vohryzek et al., 2020). The outer product of *V_1_(t)* reveals the FC matrix associated with the leading eigenvector at time *t*.

#### 2.8.2 Mode extraction

To identify recurring spatiotemporal modes *ψ* or phase-locking patterns, we clustered the leading eigenvectors for each run with K-means clustering with 300 replications and up to 400 iterations for 2-7 centroids considering 116 and 90 (i.e., excluding the cerebellum) anatomical regions. K-means clustering returns a set of K central vectors or centroids in the form of *Nx1* vectors *V_c_*. As *V_c_* is a mean derived variable, it may not occur in any individual subject data set. To obtain time courses related to the extracted modes at each TR we assign the cluster number to which *V_c_(t)* is most similar using the cosine distance.

#### 2.8.3 Mode visualization

We rendered the centroid vectors *V_c_* in cortical space by representing each element as a sphere placed at the center of gravity of the relevant brain region, and scaling the color of the spheres according to the value of the relevant eigenvector. Regions with similar phase orientation are colored alike (yellow-to-red for the smallest community and cyan-to-blue for the largest community), where darker colors (red/blue) indicate weak contributions and lighter colors (cyan/yellow) indicate stronger contributions. We also plot links between the corresponding areas to highlight the network formed by the smallest community of brain areas.

#### 2.8.4 Cluster representation in voxel space

To obtain a visualization in voxel space of the spatial modes *V_c_* we first reduced the spatial resolution of all fMRI volumes from 2mm^3^ to 10mm^3^ to obtain a reduced number of brain voxels (here *N* = 1821) to be able to compute the eigenvectors of the *NxN* phase-locking matrices. The analytic signal of each 10mm^3^ voxel was computed using the Hilbert transform, and the leading eigenvectors were obtained at each time point (with size *NxT*). Subsequently, the eigenvectors were averaged across all time instances assigned to a particular cluster, obtaining in this way, for each cluster, a *1xN* vector representative of the mean phase-locking pattern captured in voxel space.

### 2.9 Measures and metrics

The following sections provide an accessible overview of the measures and metrics used in this study. Detailed mathematical treatment and explanations for all metrics may be found in Supplementary methods and metrics. Each metric has found application in either theoretical or empirical studies, or both. Examples of their application may be found in Table 1.

**Table 1.**
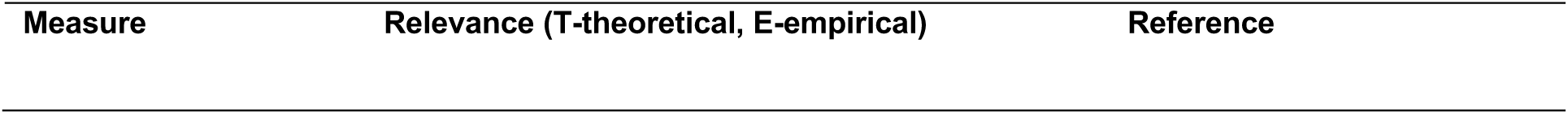
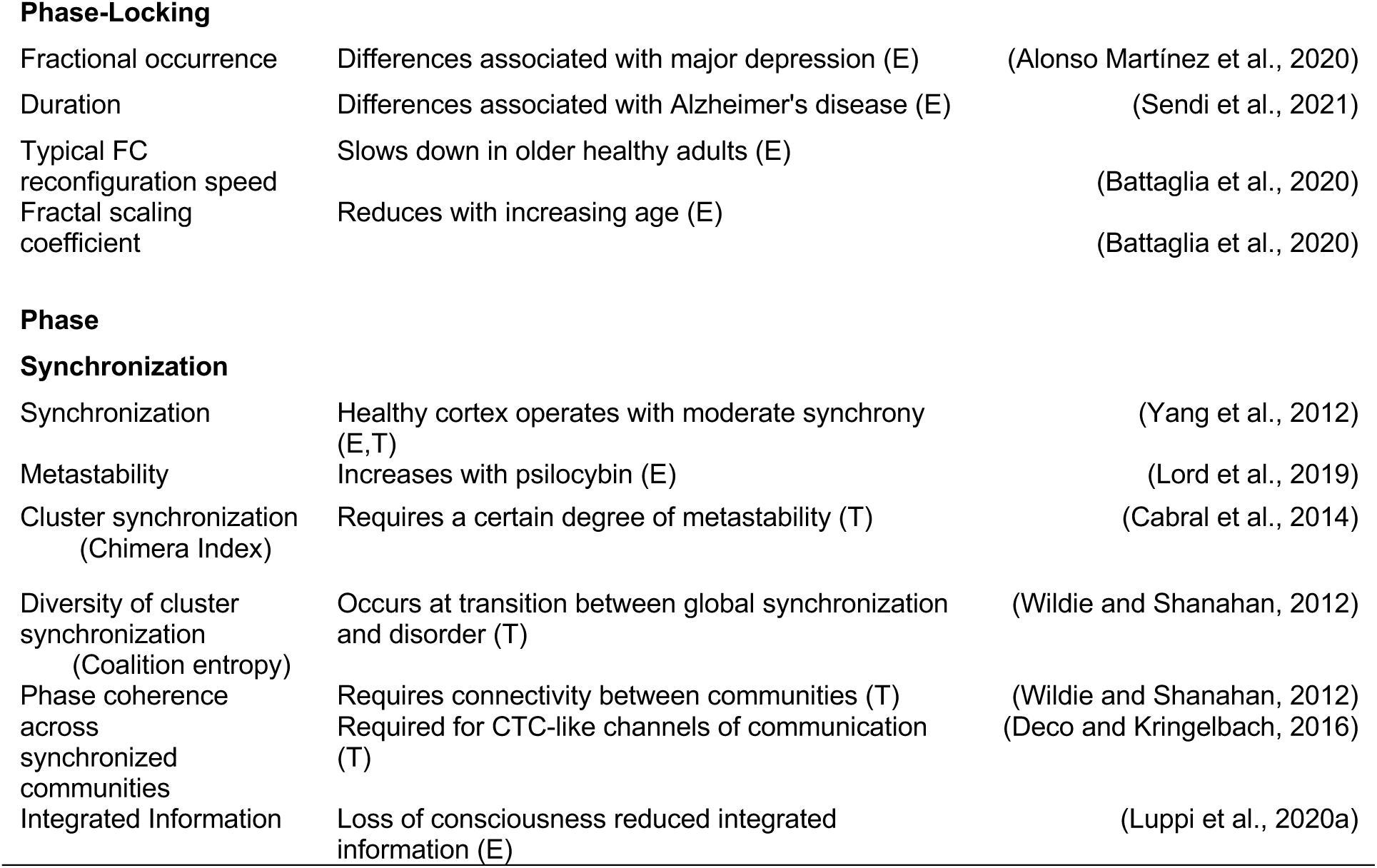
Examples of application of dFC metrics in empirical and theoretical studies

| Measure | Relevance (T-theoretical, E-empirical) | Reference |
| --- | --- | --- |
| Phase-Locking |  |  |
| Fractional occurrence | Differences associated with major depression (E) | (Alonso Martínez et al., 2020) |
| Duration | Differences associated with Alzheimer's disease (E) | (Sendi et al., 2021) |
| Typical FC reconfiguration speed | Slows down in older healthy adults (E) | (Battaglia et al., 2020) |
| Fractal scaling coefficient | Reduces with increasing age (E) | (Battaglia et al., 2020) |

|  |  |  |
| --- | --- | --- |
| Synchronization | Healthy cortex operates with moderate synchrony (E,T) | (Yang et al., 2012) |
| Metastability | Increases with psilocybin (E) | (Lord et al., 2019) |
| Cluster synchronization (Chimera Index) | Requires a certain degree of metastability (T) | (Cabral et al., 2014) |
| Diversity of cluster synchronization (Coalition entropy) | Occurs at transition between global synchronization and disorder (T) | (Wildie and Shanahan, 2012) |
| Phase coherence across synchronized communities | Requires connectivity between communities (T)<br>Required for CTC-like channels of communication (T) | (Wildie and Shanahan, 2012)<br>(Deco and Kringelbach, 2016) |
| Integrated Information | Loss of consciousness reduced integrated information (E) | (Luppi et al., 2020a) |

#### 2.9.1 Metrics derived from phase-locking

Fractional occurrence of mode *ψ*_*k*_ was calculated as number of timepoints assigned to mode *ψ*_*k*_ divided by the total number of timepoints. This measure reflects the proportion of time the fMRI activity patterns are closer to mode *ψ*_*k*_ than to any other mode *ψ*_≠*k*_. Its values are bound between 0 and 1.

Duration of mode *ψ*_*k*_ was calculated as the mean of all consecutive periods spent in a particular mode measured in seconds.

Reconfiguration speeds were calculated as 1 – correlation between functional connectivity (*iPL* matrices) at time *t* and *t+1.* This measure characterizes the time evolution of the phase-locking modes. Low speed indicates smooth transitions in phase-locking relationships. Faster speed indicates abrupt switching between phase-locking relationships.

The detrended fluctuation analysis exponent *α* returns an estimate of how predictable a timeseries is by quantifying the dependence of a value at time *t* is on a value at time *t-1*. Values less than 0.5 indicate non-persistent fluctuations and a return to the mean. Values = 0.5 indicate random fluctuations and an underlying process with no memory. Values between 0.5 and 1 indicate persistent fluctuations and an underlying process that has memory and long-term correlations.

Following (Ton and Daffertshofer, 2016), power-law scaling was tested for linearity using a Bayesian model comparison technique and the best fit model was selected with Bayesian Information Criterion. Only subjects that exhibited extended linear power-law scaling were included in the summary metric of DFA_*α*_.

#### 2.9.2 Metrics derived from phase synchrony

Communities were defined as a set of regions that intermittently lock out of phase with the global mode. The global mode was also considered as a community yielding 5 communities in total.

Synchronization was calculated as the time-average of the Kuramoto order parameter in each community, which is given by

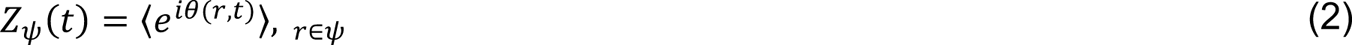

Above, *Z*_ψ_(*t*) is a complex value where its magnitude, and hence *SYNC*_ψ_= |*Z*_ψ_(*t*)|, provides a quantification of the degree of synchronization of the community at each time *t,* taking values between 1 (for fully synchronized systems) and 0 (for fully random systems.

Metastability was calculated as the standard deviation of the Kuramoto order parameter in each community. The mean value of this measure across communities denoted as global metastability, represents the overall variability in the synchronization across communities.

Cluster synchronization was calculated as the variance over communities of the Kuramoto order parameter at time t. This metric reveals if some communities cluster together in synchrony whilst other communities remain disordered.

Instantaneous phase coherence across communities was calculated as the average phase across communities with synchronization values higher than a synchronization threshold *λ*> 0.8 at time *t*. This measure represents the coherence between communities when they are highly synchronized internally. Phase coherence coefficient was calculated as the fraction of time that instantaneous phase coherence occurred.

Coalition entropy was calculated as the entropy of the coalitions formed at each timepoint *t* reported in bits. This metric represents the diversity of cluster synchronization.

Integrated information Φ^*R*^(Mediano et al., 2021) was computed from 5 binarized time-series, one for each mode extracted through K-means clustering. Values were set to 1 if synchronization values were higher than a synchronization threshold *λ* > 0.8 at time *t*. In this study, integrated information Indicates the degree of synergistic and transfer information processing within the system computed over an integration timescale τ reported in bits.

### 2.10 Statistical analysis

#### 2.10.1 Interclass correlation coefficient (ICC)

ICC is a relative metric that is used for test-retest reliability in measurement theory. It is generally defined as the proportion of the total measured variance that can be attributed to within subject variation. As such, ICC coefficients may be low when there is little variance between subjects, that is in a homogeneous sample, or when the within-subject variance is large (Xing and Zuo, 2018).

There are many scales for ICC, so for clarity we will use those of (Landis and Koch, 1977):

- low (0 < ICC < 0.2)
- fair (0.2 < ICC < 0.4)
- moderate (0.4 < ICC < 0.6
- substantial (0.6 < ICC < 0.8)
- almost perfect (0.8 < ICC < 1)

We calculated the run reliability of mode *ψ* extraction with ICC(1,1) in search of agreement rather than consistency across runs (Noble et al., 2021). For the test-retest assessment of metric consistency over runs, we used the ICC(3,1) form (Shrout and Fleiss, 1979) as recommended by (Koo and Li, 2016) which is the equivalent of a 2-way mixed ANOVA. As such, there is an assumption that the data comes from a normal distribution. When the assumption of normality is violated, it is recommended to use non-parametric tests such as permutation testing.

#### 2.10.2 Repeated measures ANOVA

We performed repeated measures ANOVA on global metrics using the *ranova()* function in MATLAB MathWorks R2021b. Greenhouse-Geisser correction was necessary as the assumption of sphericity was violated in most cases. We therefore assessed normality of the data with Chi-square goodness of fit (results not included). As the results indicated non-normal distribution of the data, we decided to replace ICC(3,1) with non-parametric permutation testing. We also performed repeated measures ANOVA on the mode-specific metrics. It should be noted that for the AAL parcellation that included the cerebellar regions, the order of the modes in run 2 and 4 was adjusted to match the order in run 1 and 3 for all statistical testing.

#### 2.10.3 Permutation testing

We used a non-parametric permutation-based paired t-test to identify significant differences between runs. This non-parametric two-sample hypothesis test uses permutations of group (run) labels to estimate the null distribution rather than relying on the t-test standard distributions. The null distribution was computed independently for each run. A t-test was then applied with 1000 permutations to compare runs.

#### 2.10.4 Linear mixed effects modeling

We used lmerTest (Kuznetsova et al., 2017) in RStudio 2021.09.1 Build 372, with the purpose of building predictive models with both standardized and non-standardized metric data that could deal with data that was not independent and identically distributed. To investigate the relationship between integrated information and all other metrics, we fitted a linear mixed-effect model (estimated using REML and nloptwrap optimizer) to predict PHI with standardized metric values. The model included RUN as random effect (formula: ∼1 | RUN). 95% Confidence Intervals (CIs) and p-values were computed using the Satterthwaite’s method.

### 2.11 Code availability statement

The Matlab and R code developed for this analysis will be made available on publication at github.com/franhancock/Complexity-science-in-dFC together with the 5 phase-locking mode centroids for AAL parcellation in NIFTI and in Matlab format.

## 3 Results

### 3.1 Reliability of dFC measures and metrics

#### 3.1.1 Spatial patterns of phase-locking are invariant across fMRI acquisitions

We first sought to evaluate if the spatiotemporal patterns of phase-locking observed in fMRI are representative and stable across multiple acquisitions. For this purpose, we compared the spatial patterns of phase-locking extracted independently for each of the 4 fMRI runs recorded from the same 99 participants (Figure 2). Each mode of phase locking *ψ*_*k*_ corresponds to a 1xN vector (with N being the number of brain areas considered) obtained through K-means clustering of phase-locking patterns obtained at every time point in each run. We chose K=5 modes considering previous test-retest studies (Abrol et al., 2017; Vohryzek et al., 2020). We calculated run reliability with inter-class correlation coefficient (ICC) in search of agreement across runs (Noble et al., 2021). With N=90 anatomical non-cerebellar brain regions defined in the AAL parcellation, the modes extracted independently in each run showed almost perfect agreement between runs with 0.99 > ICC > 0.97. With the inclusion of cerebellar regions, the reliability of spatial patterns showed again almost perfect agreement between runs with 1 > ICC > 0.94, although the probability of occurrence differed across runs, altering the order of the modes when sorted by relative occupancy.

**Figure 2.**
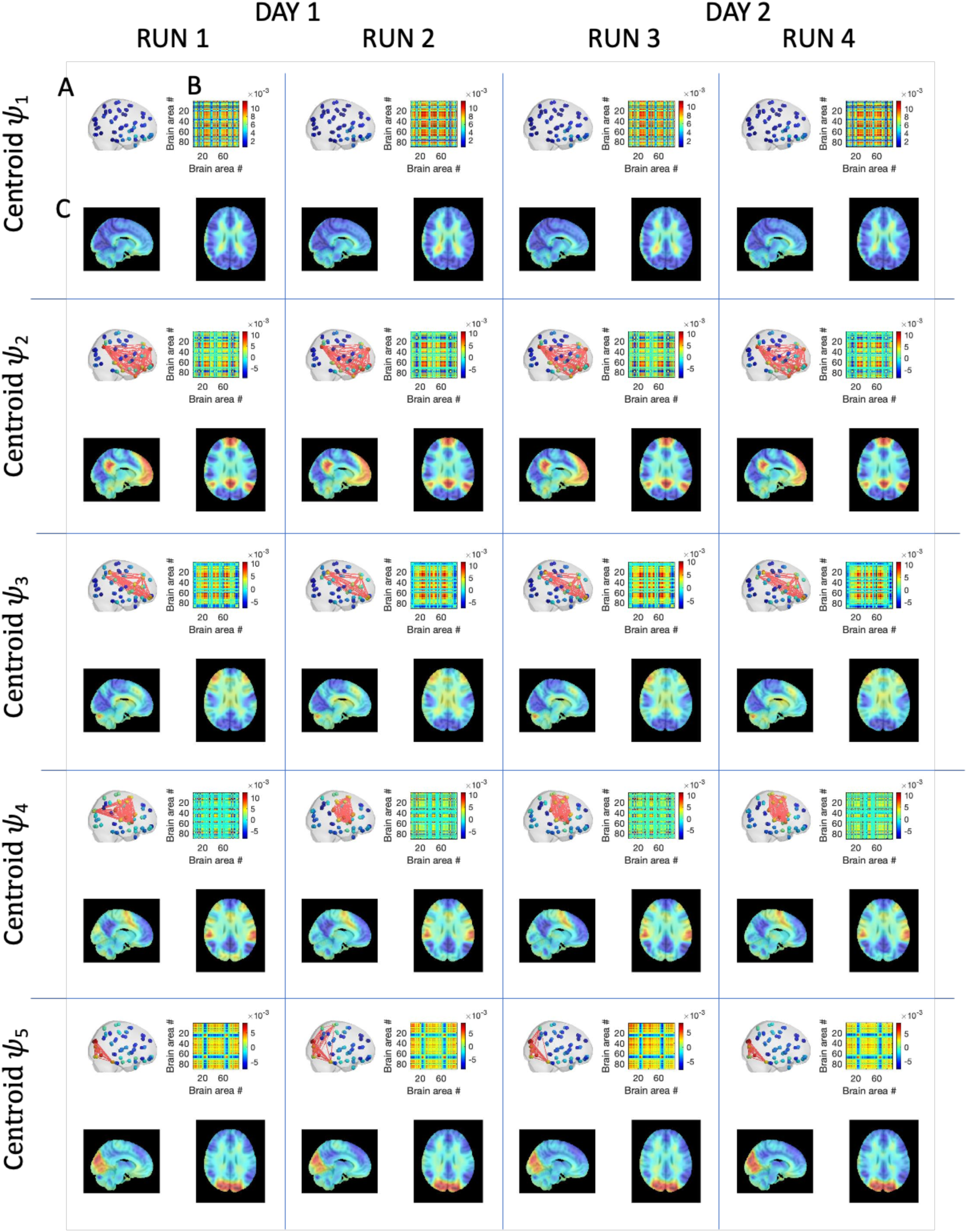
- Invariant spatiotemporal patterns of phase-locking obtained independently in each of the 4 fMRI runs on the same 99 participants. LEiDA was applied separately to the 4 fMRI runs recorded on 2 consecutive days from 99 participants and the centroids obtained from clustering into K=5 are reported here. Each centroid V_c_ (with size 1xN, with N=90) is represented in three distinct forms: (A) each element V_c_(n) is represented as a sphere placed at the center of gravity of the corresponding brain region and its color is scaled according to its value in V_c_. Links highlight the network formed by the smallest community of brain areas. (B) the phase-locking matrices computed as the outer product of the centroid vector V_c_. (C) Representation of the centroid vector for each mode in 10mm voxel space by averaging the eigenvector values over all time instances assigned to a particular cluster/mode. The modes were then plotted over a 1mm^3^ MNI T1 image.

The similarity of the 5 cluster centroids *ψ*_*K*=1,..,5_ across the 4 runs is clearly visible in Figure 2. To illustrate the patterns of phase relationships between brain regions, the 1xN centroids are rendered in cortical space together with the associated phase-locking matrices. In addition, to visualize the phase-relationships in voxel space, we reduce the fMRI volumes from 2mm^3^ to 10mm^3^, resulting in 1821 brain voxels within the MNI brain mask, and compute the eigenvectors of phase-locking at each time point. Subsequently, the eigenvectors are averaged across all time points assigned to each cluster, and represented in sagittal and axial planes overlaying on a 1mm^3^ MNI structural image. This approach allows visualizing the patterns of phase relationships in voxel space, revealing meaningful functional subsystems overlapping with resting-state networks described in the literature (A similar figure when 116 regions are considered in the K- means clustering can be found in inline Supplementary Figure 1. ICC values for both 90 and 116 regions are reported in inline Supplementary Figure 2).

We used the eigenvectors obtained with 116 regions to shed light on the composition of the extracted modes and their putative membership of seven cerebral intrinsic functional networks (Yeo et al., 2011) collectively known as resting-state networks (RSN), and connections with the sub-cortical and cerebellum regions. In inline Supplementary Figure 3 we show the composition of each mode eigenvector color-coded according to the RSNs, and the rendering of these eigenvectors in cortical space.

We find that mode *ψ*_1_ represents a global mode where the fMRI signals in all regions are aligned in-phase. Mode *ψ*_2_ consists of a phase-locking pattern where regions associated with the Default Mode Network (DMN), the Limbic network (LBC), the subcortical hippocampi (SC) regions, and some cerebellum (CB) regions are shifted in phase with respect to the rest of the brain. Mode *ψ*_3_ comprises regions associated with the Frontal Parietal Area (FPA), the LBC, the SC Caudate and Putamen, and a number of CB regions. Mode *ψ*_4_ comprises of regions associated with the Sensory Motor network (SMT), and the Ventral Attention network (VAT), with some contribution from the FPA and the CB regions. Finally, *ψ*_5_ is comprised mainly of the Visual network (VIS) with significantly lower contributions from SMT, LBC, DMN, and SC.

Overall, these results show that spatiotemporal patterns of phase-locking are representative and stable across multiple fMRI acquisitions. They therefore provide a stable basis for the characterization and analysis of our battery of dFC metrics.

#### 3.1.2 Global Metastability was the most stable metric across all runs

As a second step, we sought to investigate the stability of a series of global metrics - namely metastability, synchronization, chimera index, phase-coherence coefficient, coalition entropy, integrated information, and typical reconfiguration speed – across different multiple fMRI acquisitions. For this, the values of each metric in four different runs were compared using a non-parametric permutation-based paired t-test to identify significant difference. Figure 3A shows the bar plots for each metric including the mean value and indicators for where significant differences were found between the runs. In Figure 3B we show the distribution of the metrics across runs which provides complementary information on the median and spread of the metric values across runs.

**Figure 3.**
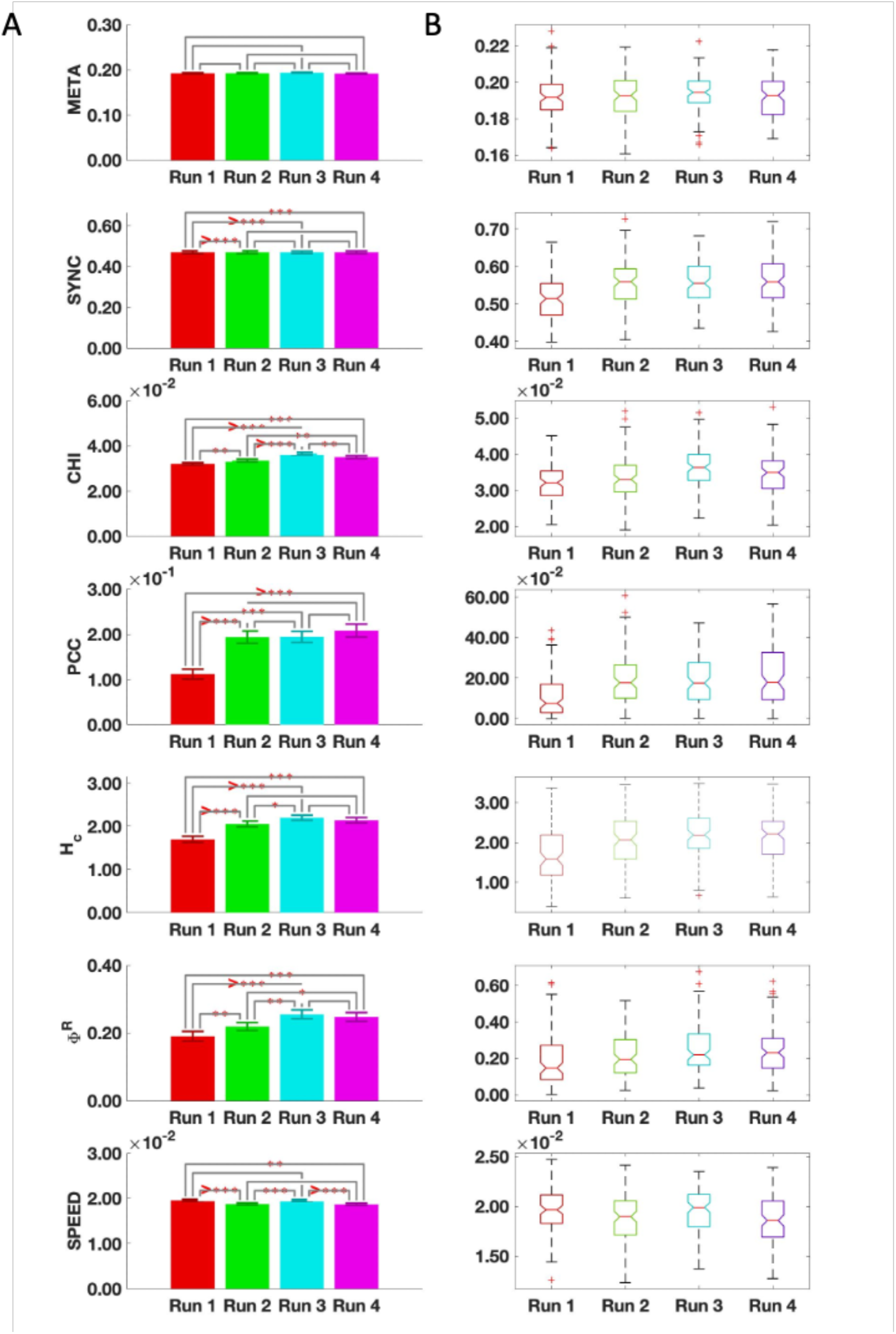
Stability of global metrics across 4 runs. (A) The mean values for each metric are shown as bar plots. The *** indicate a statistically significant difference between the metric across the associated runs where * *p* < 0.05, ** *p* < 0.01, *** *p* < 0.001, and >*** *p* < 0.0001. (B) The distribution of the global metrics across runs. META, metastability; SYNC, synchronization; CHI, chimera index; PCC, phase-coherence coefficient; HC coalition entropy; Φ^*R*^, integrated information; SPEED, typical reconfiguration speed.

There was no statistically significant difference in the measure of global metastability across the 4 fMRI runs. When the cerebellum was excluded, however, global metastability did not show the same reliability inline Supplementary Figure 4(A-B). The measures of global synchronization and phase-coherence coefficient were found to be reliable across runs 2, 3, and 4. The remaining metrics however, showed statistically significant differences across the 4 acquisitions. To test the contribution of the cerebellar regions for the reliability of the metrics, we performed the same analysis with NEUROMARK (Du et al., 2020), a parcellation based on intrinsic connectivity networks that includes these regions. Indeed, in this case, global metastability remained representative and stable across all 4 runs inline Supplementary Figure 4(C-D).

#### 3.1.3 High dynamical- and informational-complexity across acquisitions of resting-state fMRI

Although a measure of global metastability was found to be stable across the cohort of healthy young adults between all runs, this was not the case for individual subjects. To illustrate this, we plot the temporal evolution of a series of metrics for two scans from one representative subject as illustrated in Figure 4. For comparison purposes, we include the same information for another subject in inline Supplementary Figure 5.

**Figure 4.**
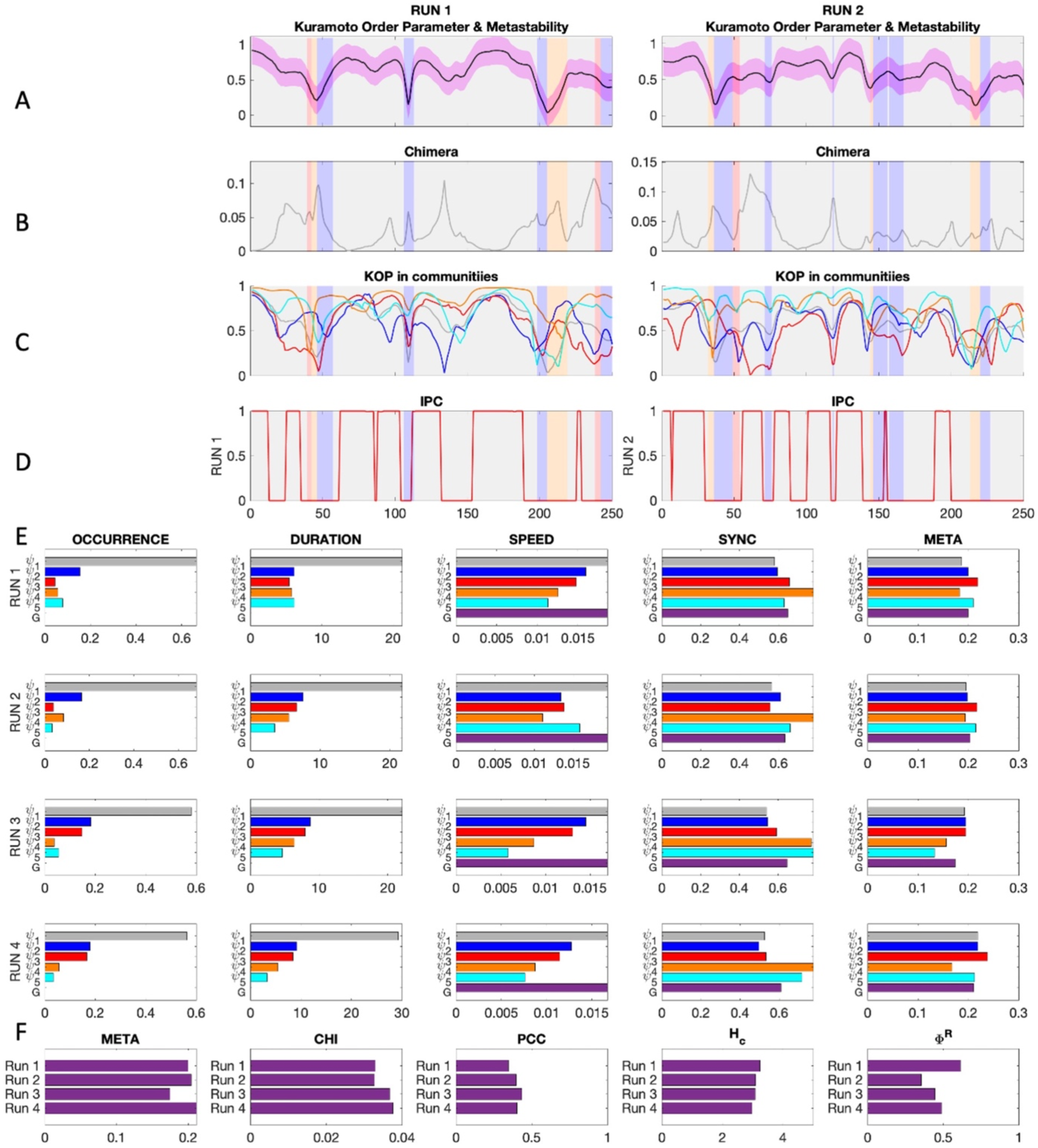
Overview of all metrics in all runs for a representative subject. (A) Exemplar snippets from the instantaneous phase synchrony or Kuramoto order parameter time series for each run color-coded to show which mode was dominant over time. (B) The same as A but for chimeras or cluster synchronization. (C) The evolution of instantaneous synchrony within each of the color-coded modes. (D) The evolution of instantaneous phase-coherence. (E) Mode-specific metrics calculated independently for each of the 4 runs. (F) The values of the global metrics across all 4 runs. META, metastability; CHI, chimera index; PCC, phase coherence coefficient; H_c_, coalition entropy; and Φ^*R*^, integrated information.

#### 3.1.4 Mode-specific metrics do not appear representative or stable across runs

We further defined mode-specific metrics by considering only the subsets of brain areas shifted in phase in each spatial mode, and compared their values across the 4 runs. Mode-specific metrics are commonly used to investigate differences between normal and abnormal functional brain activity (Kottaram et al., 2019; Zarghami et al., 2020). Using repeated measures ANOVA, we did not find any mode-specific metric that was reliable in all 5 modes across all 4 runs when excluding or including the cerebellar region as shown in Table 2.

**Table 2.** Repeated measures ANOVA results for mode-specific metrics over 4 fMRI acquisitions in AAL parcellations excluding the cerebellar regions (AAL90), and including the cerebellar regions (AAL116). OCC, occurrence; META, metastability; SYNC, synchronization; SPEED, typical reconfiguration speed.

| Repeated measures ANOVA for mode-specific metrics<br>Significant differences<br>AAL90 |  |  |  | Repeated measures ANOVA for mode-specific metrics<br>Significant differences<br>AAL116 |  |  |  |
| --- | --- | --- | --- | --- | --- | --- | --- |
| Metric | Mode | F score | p value | Metric | Mode | F score | p value |
| OCC | $\psi_2$ | $F(3,294) = 10.570$ | $p < 0.001$ | OCC | $\psi_1$ | $F(3,294) = 7.158$ | $p < 0.001$ |
| OCC | $\psi_4$ | $F(3,294) = 5.362$ | $p = 0.001$ | OCC | $\psi_2$ | $F(3,294) = 9.634$ | $p = 0.001$ |
| DURATION | $\psi_1$ | $F(3,294) = 6.139$ | $p = 0.001$ | OCC | $\psi_3$ | $F(3,294) = 2.817$ | $p = 0.039$ |
| DURATION | $\psi_2$ | $F(3,294) = 3.152$ | $p = 0.025$ | OCC | $\psi_4$ | $F(3,294) = 8.234$ | $p < 0.001$ |
| META | $\psi_5$ | $F(3,294) = 7.462$ | $p < 0.001$ | DURATION | $\psi_1$ | $F(3,294) = 7.932$ | $p < 0.001$ |
| SYNC | $\psi_1$ | $F(3,294) = 14.466$ | $p < 0.001$ | META | $\psi_3$ | $F(3,294) = 4.262$ | $p = 0.006$ |
| SYNC | $\psi_2$ | $F(3,294) = 7.062$ | $p < 0.001$ | SYNC | $\psi_1$ | $F(3,294) = 11.334$ | $p < 0.001$ |
| SYNC | $\psi_3$ | $F(3,294) = 9.381$ | $p < 0.001$ | SYNC | $\psi_2$ | $F(3,294) = 5.138$ | $p = 0.002$ |
| SYNC | $\psi_4$ | $F(3,294) = 24.355$ | $p < 0.001$ | SYNC | $\psi_3$ | $F(3,294) = 3.537$ | $p = 0.015$ |
| SYNC | $\psi_5$ | $F(3,294) = 6.956$ | $p < 0.001$ | SYNC | $\psi_4$ | $F(3,294) = 17.132$ | $p < 0.001$ |
| SPEED | $\psi_1$ | $F(3,294) = 9.069$ | $p < 0.001$ | SYNC | $\psi_5$ | $F(3,294) = 85.632$ | $p < 0.001$ |
| SPEED | $\psi_2$ | $F(3,294) = 12.065$ | $p < 0.001$ | SPEED | $\psi_1$ | $F(3,294) = 8.552$ | $p < 0.001$ |
| SPEED | $\psi_3$ | $F(3,294) = 9.251$ | $p < 0.001$ | SPEED | $\psi_2$ | $F(3,294) = 16.348$ | $p < 0.001$ |
| SPEED | $\psi_4$ | $F(3,294) = 8.907$ | $p < 0.001$ | SPEED | $\psi_3$ | $F(3,294) = 3.272$ | $p = 0.022$ |
| SPEED | $\psi_5$ | $F(3,294) = 10.805$ | $p < 0.001$ | SPEED | $\psi_4$ | $F(3,294) = 17.662$ | $p < 0.001$ |
| | | | | SPEED | $\psi_5$ | $F(3,294) = 65.839$ | $p < 0.001$ |

We also used a non-parametric permutation-based paired t-test to investigate if these differences were due to 1 idiosyncratic run or if the differences emerged over different runs. The differences remained statistically significant and were present across different runs even after performing Bonferroni correction for multiple comparisons across the 5 modes, as can be seen in inline Supplementary Figure 6. Notably, metrics of fractional occurrence and duration of modes - which have been used for comparisons between conditions studies - were not reliable across 4 acquisitions in the AAL parcellation with or without the cerebellar regions.

### 3.2 Characterization of the dFC process

#### 3.2.1 Reconfiguration speeds in phase-locking space exhibit fractal scaling and deviate from Gaussianity

An unanswered question regarding dFC is whether spatiotemporal patterns change in a discrete or continuous manner over time. K-means clustering yields a distinct mode for each timepoint, but this mode is just the cluster centroid with the shortest distance, and a number of other modes may also contribute to the resulting spatiotemporal pattern at each timepoint. An alternative perspective is to view dFC as a smooth reconfiguration of phase-locking connectivity, and to collapse these relations to a point in the space of possible relations. We can then view the evolution of this point as a stochastic exploration of a high-dimensional space. This is a direct adaptation of the reconfiguration speed introduced in (Battaglia et al., 2020) for phase-locking functional connectivity.

We computed the reconfiguration speeds (Figure 5A) and fractal scaling characteristics (Figure 5D) of phase-locking dFC. Out initial plan was to include the Hurst-like exponent *α* derived from detrended fluctuation analysis (DFA_*α*_) (Peng C-K et al., 1993) in our battery of dFC metrics (see 2 Materials and Methods). However, we found that the assumption of extended linear power-law scaling was violated in 40-50% of subjects Figure 5(B-C). When linear power-law scaling was present, FC fluctuations showed fractal scaling with DFA_*α*_ > 0.5 indicating that the stochastic reconfiguration process in phase-locking space was not random, but displayed long-range correlations and deviated from Gaussianity as shown in Figure 5D.

**Figure 5.**
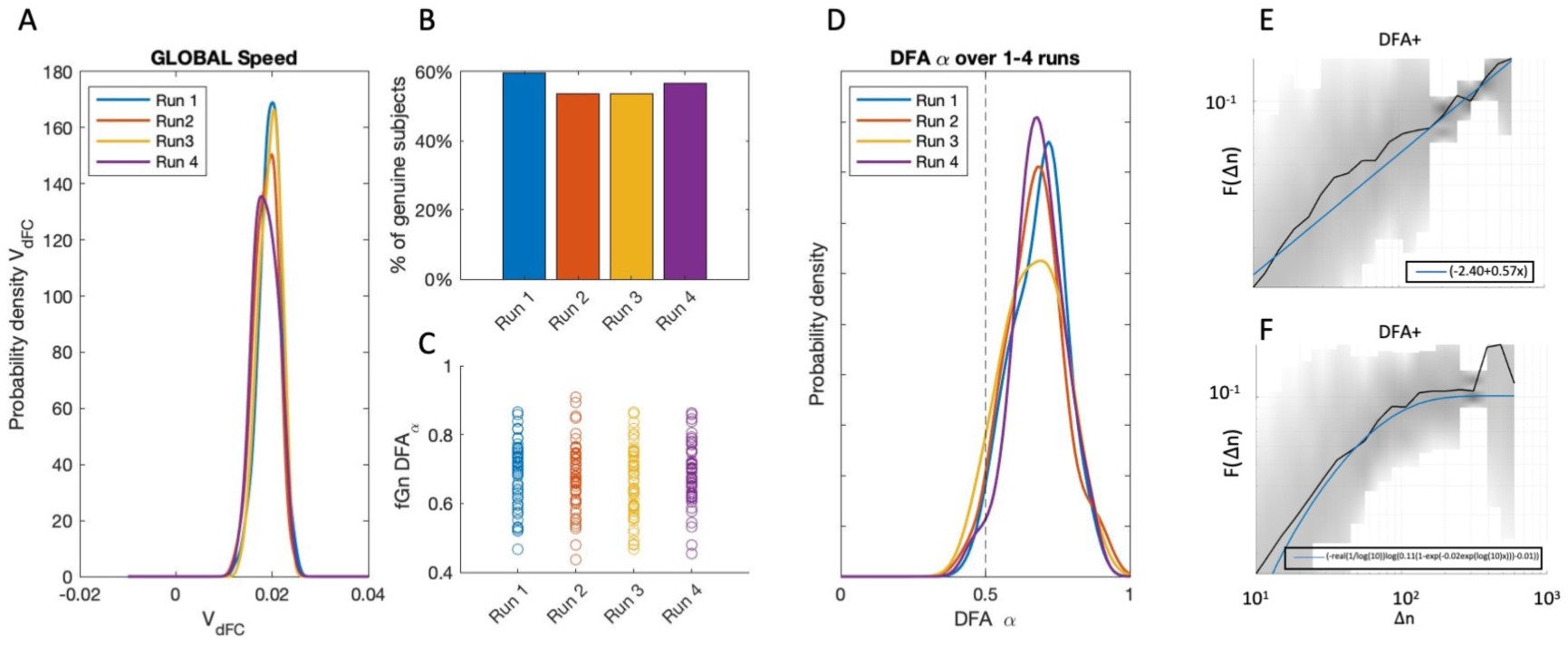
dFC reconfiguration speeds and Detrended Fluctuation Analysis (DFA). (A) Phase-coupling dFC reconfiguration speeds were slow across all 4 fMRI acquisitions. (B) Before performing DFA we established if subjects exhibited extended sections of linear power-law scaling in FC fluctuations. Between 50% to 60% of subjects exhibited ‘genuine’ power-law scaling. (C) For the majority of subjects that demonstrated linear power-law scaling, *DFA_α_* was greater than 0.5 which implies the presence of persistent fluctuations, long-range correlations, and deviation from a Gaussian generation process. (D) The probability densities for *DFA_α_* for ‘genuine’ subjects in each of the 4 runs. (E) An example of linear power-law scaling where the best fit was found to be y=2.40*0.57x. (F) An example of non-linear power-law scaling where the best fit was found to be y= (-real(1/log(10))log(0.11(1-exp(- 0.02exp(log(10)x)))-0.01)). We used FluctuationAnalysis() (Ton and Daffertshofer, 2016) to test for linearity and to calculate *DFA*.

The reconfiguration random-walk of dynamic phase-locking matrices, or *dPL stream*, is represented in 3 dimensions in Figure 6 (using a t-Stochastic Neighbor Embedding algorithm, see Materials and Methods). The resulting distance preserving non-linear projections in 3 dimensions of the associated *dPL stream* (timeseries) are shown with respect to time (left) and with respect to the mode visited (right). The speeds of reconfiguration revealed periods of slow morphing interspersed with sharp changes in the configuration of phase-locked connectivity corresponding to the concept of ‘knots and leaps’ in (Battaglia et al., 2020), in contrast to unstructured space filling as would be expected for uncorrelated speeds in a memoryless stochastic process..

**Figure 6.**
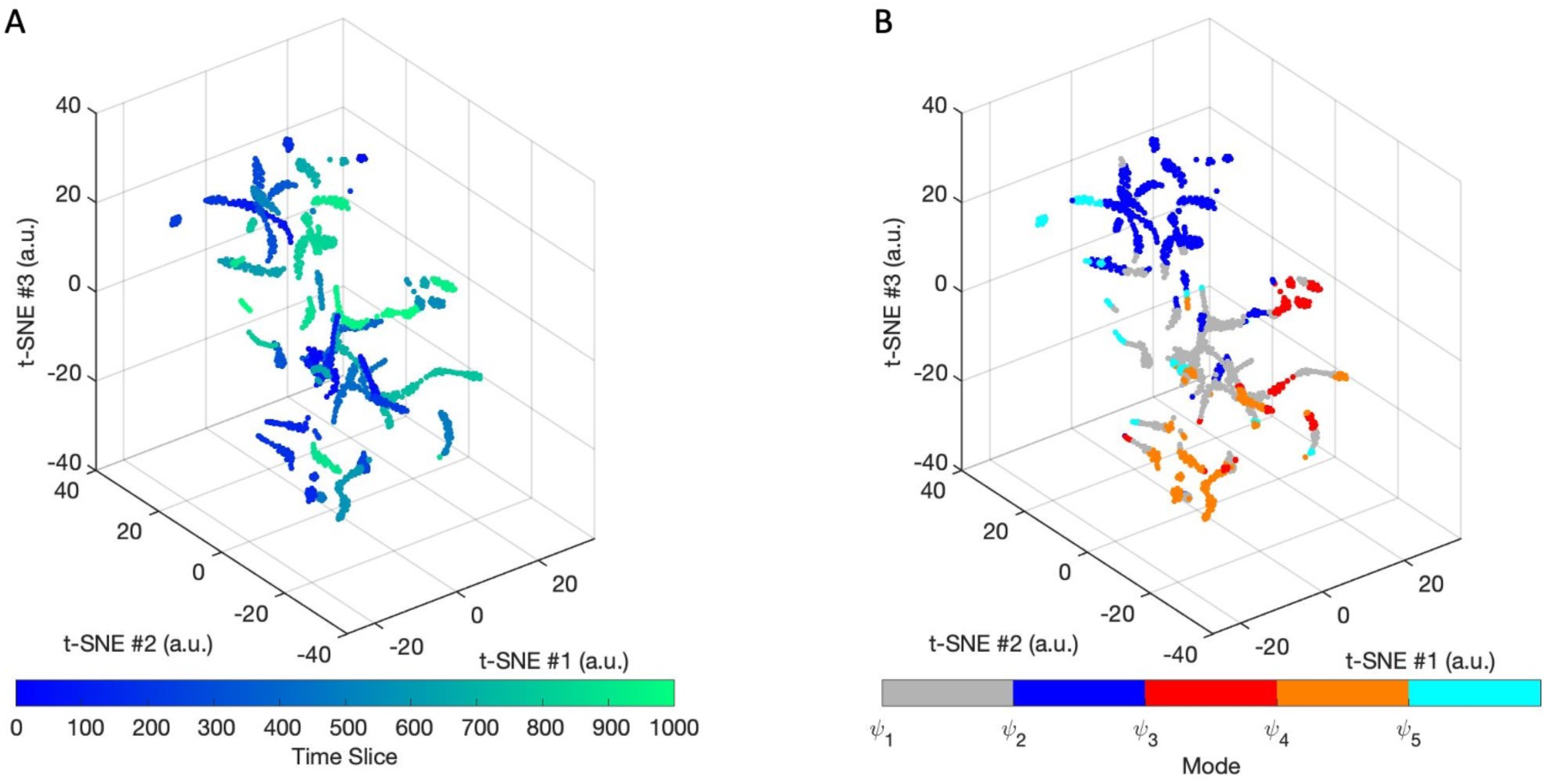
Visualizations of a reconfiguration walk in the phase space of leading eigenvectors. (A) We show a distance preserving non-linear projection in three dimensions of a subject’s *dPL* stream from a single fMRI scan obtained with the t-SNE algorithm. Each point corresponds to a specific observation of FC(t) and the path connecting the points indicates in (A) the temporal order in which the different configurations are visited. (B) The same projection but color-coded with the mode assigned to the timepoint in the timeseries.

Overall, these findings are consistent with previous literature (Battaglia et al., 2020) and suggest that spatiotemporal patterns of phase-locking change in a non-random slow continuous fashion over time.

### 3.3 Relationship between global metrics

As a next step in our investigation, we sought to investigate how the various studied global metrics are related to each other between subjects. For this, we calculated the Spearman correlation between all pairs of metrics. As illustrated in Figure 7 (which corresponds to RUN1), most metrics were significantly correlated, with some metrics correlating more than 90% - revealing relationships that can be more or less evident given their nature. For instance, it is not surprising that synchrony is highly correlated (r=0.84) with the occupancy of the mode 1, since the latter represents more time in a mode of global phase coherence, while being also highly correlated (r=0.92) with the phase coherence coefficient. Moreover, the occupancy and duration of mode 1 are also highly correlated (r=0.92), which can be explained by the fact that the more a mode occurs, the more probable it is to be detected on 2 consecutive time points. Less obvious, perhaps, are the strong correlations detected between Phase Coherence Coefficient, Coalition Entropy and Integrated Information. Moreover, both the Chimera Index (CHI) and the reconfiguration speed (SPEED) exhibit negative relationships with the other metrics, but the two are not correlated to each other, indicating that they are sensitive to complementary dynamical features of the system. Correlation matrices for runs 2-4 may be found in inline Supplementary Figure 7.

**Figure 7.**
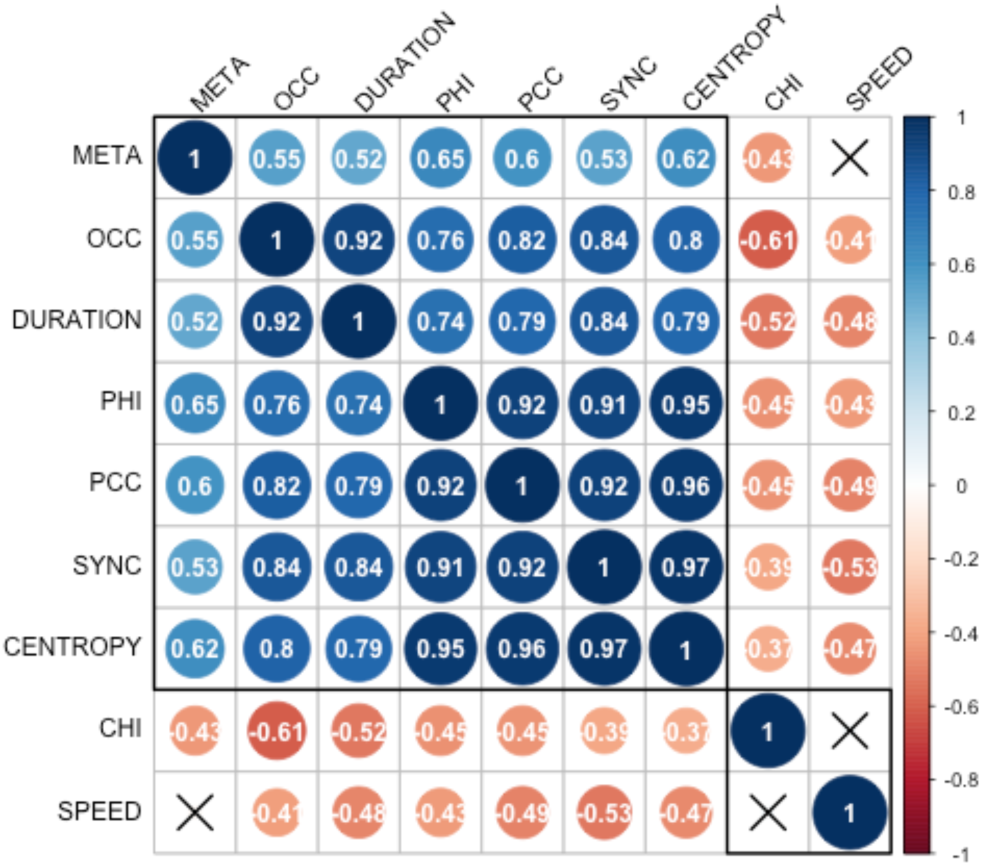
Relationships between metrics. Correlation coefficients for all metrics in run 1. Coefficients with X indicate statistical significance with *α* < 0.05. SYNC, synchronization, CHI, chimera index, META, metastability, OCC, fractional occurrence of *ψ*_1_, DURATION, duration of *ψ*_1_, SPEED, typical reconfiguration speed, PCC, phase-coherence coefficient, CENTROPY, coalition entropy, PHI, integrated information.

To further investigate the relationship between integrated information and all other metrics, we fitted a linear mixed-effect model to predict PHI based on the values of SYNC, CENTROPY, and CHI with standardized metric values. As there appeared to be quadratic structure in the distribution of the residuals, we investigated each predictor variable in its quadratic form. SYNC^2^ provided the best model fit and so was retained as a quadratic term. Additionally, the model included random intercepts to account for the effect of different fMRI runs. The model’s explanatory power related to the fixed effects alone (i.e. its marginal R^2^) is 0.85. All predictors in this model were found to be significantly correlated, with SYNC and CENTROPY having positive effects while CENTROPY having a negative effect as illustrated in Table 3.

**Table 3.** Linear mixed-effect regression model - fixed effects

| Predictor | Beta | T score | p value |
| --- | --- | --- | --- |
| SYNC<sup>2</sup> | 0.05 | t(390) = 3.03 | p = 0.003 |
| CENTROPY | 0.87 | t(390) = 42.35 | p < 0.001 |
| CHI | -0.11 | t(390) = -5.17 | p < 0.001 |

The quality of the model fit was assessed using *performance* (Lüdecke et al., 2021) and a visualization of the model checks can be found in inline Supplementary Figure 8. Additionally, this linear mixed regression model indicated that there were no random effects due to RUN as the standard deviation of the random intercept was 1.454e-16.

These findings provide unique empirical evidence that dynamical- and informational-complexity are related; and shows convergence of evidence from multiple approaches to support the interpretability of these metrics with respect to neuroscience.

## 4 Discussion

When conceptualizing the brain as a complex system (Turkheimer et al., 2021) one has a number of theoretical approaches and corresponding methodological tools available to assess dynamic functional connectivity beyond viewing them as mere time-varying temporal correlations in fMRI signals. In this work we empirically investigated the relationships between approaches that investigate intrinsic brain activity from a dynamical systems perspective, from a stochastic process view, and from an information-processing perspective, providing some practical first steps towards the development of unified accounts of brain function.

Four main insights can be derived from our results. First, from a methodological perspective, phase-locking functional connectivity derived with LEiDA, provides an invariant basis of spatial modes for the investigation of dynamical behavior between brain regions. This invariant basis could be used as a template for future studies providing a validated (in terms of test-retest reliability) basis for cross study comparisons. These 5 reliable spatiotemporal modes of phase-locking activity reflect the physics of self-organization (Haken, 1996): that is, these macroscopic patterned modes are spontaneously created and change dramatically at critical points, showing how global order can emerge from local interactions (Kelso, 1995).

Second, global metastability was the only representative and stable metric across a cohort of healthy young adults when the cerebellum is considered in conjunction with the cortex and subcortex. This may seem surprising given that the modes themselves were invariant across scanning sessions. However, the modes reflect centroids derived from k-means clustering, and as such represent the center of the cluster. Any particular instance or realization of a fMRI timeseries will not necessarily reflect these centroids, but will nevertheless have their time-points assigned to the mode they are closest to. The disadvantage of such hard clustering is that each time-point will only be assigned to one mode when in fact, the spatiotemporal pattern of the time-point may closely match more than one mode.

In addition to methodological considerations, there may be physiological effects that affect brain activity across runs. Indeed, within-individual changes in resting-state dynamics have been associated with fluctuations in arousal (Laumann et al., 2017), physiological state (Chang et al., 2013; Schneider et al., 2016), ongoing conscious experience (Gonzalez-Castillo et al., 2021) and spontaneous memory replay (Tambini and Davachi, 2019). Systematic differences have also been found with time of day (Orban et al., 2020; Vaisvilaite et al., 2021). However, our results indicate that global metastability is relatively insensitive to these effects. This global metric is therefore a potential candidate for neurological markers of effect in intervention studies.

Indeed, empirical results have shown global metastability to be higher when the brain was at rest (Hellyer et al., 2014), reduced during states of unconsciousness (Jobst et al., 2017), and increased beyond the resting-state maximum when the brain was in a psychedelic state (Carhart-Harris et al., 2014; Lord et al., 2019). In clinical populations, global metastability was found to be progressively reduced for mild cognitive impairment to Alzheimer’s disease (Córdova-Palomera et al., 2017) and positively correlated with cognitive flexibility (Hellyer et al., 2015). Metastable synchronization of brain subsystems has also been shown to drive the transient emergence of cluster synchronization, replicating features of resting-state magnetoencephalography MEG (Cabral et al., 2014). Global metastability, is therefore, a reliable dFC metric that has promise for both empirical and computational studies.

However, as the majority of metrics were not representative across the same subjects in different acquisitions, they may not be representative or generalizable to the overall population of healthy young adults. This nonergodicity challenges the interpretation of cross-sectional study outcomes and questions the applicability of such designs to study phenomena that may be more suitable to investigation of individual life-trajectories through approaches such as fingerprinting (Van De Ville et al., 2021).

The differential effect of including the cerebellum in the calculation of our dFC metrics is intriguing. The cerebellar regions have been shown to be associated with the DMN and FPA (Buckner et al., 2011), and to be active across a range of motor and cognitive tasks including working memory, cognitive control, social cognition (King et al., 2019) and emotional processing (Pierce and Péron, 2020). Interestingly, it has been suggested that the cerebellar regions fine-tune limbic-induced synchronization of the cortical regions (Pierce and Péron, 2020) which is consistent with our findings that mode *ψ*_4_ includes frontal-parietal, limbic, and cerebellar regions. This synchronization effect of the cerebellar regions has been neglected to date in dFC studies, but appears to play a key role for the reliability of global metastability.

Third, we sought to find reproducible evidence of convergence from multiple methods by investigating the relationship between our diversly derived metrics. The development of a prediction model that was independent of run and included metrics derived from dynamical systems theory, information theory, and information dynamics testifies to the neuroscientific interpretability of our results. It also revealed in empirical data that dynamical- and informational complexity are related, confirming previous computational study findings (Mediano et al., 2022). It is interesting to note that in our regression model, the main effect of cluster synchronization was to *reduce* mean integrated information Φ*^R^*. What this suggests is that excessive competition between the communities to create coalitions may lead to predominantly redundant information processing; conversely, the diversity of cluster coalitions would be what leads to transfer and synergistic information processing. Integrated information - as computed in this study, with the additional subtleties of decomposition and multivariate sources and targets, may be capturing some elements of conscious processing. Intriguingly, integrated information was not significantly predicted by metastability, although moderate positive correlations between the two metrics were found in all 4 runs. Indeed, global metastability could be associated with homeostasis, reflecting a healthy regulation of tendencies for integration and segregation, whereas integrated information theoretically reflects the actual balance of integration and segregation. Metastability can be viewed as providing the opportunity for the system to engage in cluster synchronization resulting in segregation of the communities. This dynamic segregation feature of the system appears to be complementary to the speed of changes in FC, and the metrics sensitive to these features exhibit negative relationships with all other metrics.

Taking metrics derived from dynamical systems theory and stochastic processes yields complementary insights into the dynamical complexity of brain functioning. However, not all metrics revealed findings consistent with previous literature. It is not unexpected to find periods of high phase coherence across communities in the global mode, but it would be expected to find CTC-like channels of communication when other modes are dominant. It may be that the synchronization threshold *λ* = 0.8 was too high to allow for delays in phase synchronization between remote communities. Indeed, when the threshold was set to λ = 0.7, periods of high phase coherence across communities were also found in other modes as can been seen in inline Supplementary Figure 9.

Computational models play a crucial role in neuroscience either for predicting phenomenon or for replicating phenomenon observed in empirical data. In this study we included metrics from both empirical data and computational modelling, and have unveiled relationships that will require a fundamental review of the underlying theoretical and mathematical concepts for neuroscientific interpretation.

Fourth, and finally, from describing brain behavior from the perspective of a stochastic process, we have provided tentative confirmatory results that the dFC process changes in a slow, non-random manner. It must be noted that we used phase-locking functional connectivity rather than temporal correlation as in the original application of this innovative methodology (Battaglia et al., 2020). Our non-linear measures of dynamic phase-locking behaved differently than linear correlations.

Despite this difference, we were still able to show in the majority of subjects, that spatiotemporal patterns of phase-locking change in a continuous and non-random manner, exhibiting long-range temporal correlations, indicating the presence of memory.

Taken together, these results are congruent with complex systems theory (Turkheimer et al., 2021) in that phase-relationships in fMRI of the resting state brain exhibit:

- Invariant spatiotemporal patterns that are indicative of self-organized processes (Haken, 1996)
- Nonergodicity in that dFC metrics are in general, not representative across samples (Turkheimer et al., 2021)
- Diversity in cluster synchronization (Turkheimer et al., 2021)
- Fractal scaling in the continuous change of functional connectivity (Battaglia et al., 2020)

## 5 Limitations and future research

A number of limitations that should be considered when evaluating the findings.

Starting with modes, we found near perfect ICC agreement of all 5 spatiotemporal phase-locking modes across all 4 runs. However, ICC is a relative metric and the large between-region differences may bias a high ICC value in the absence of genuinely small within-region differences. However, we achieved similar results with Pearson correlation.

Another possible limitation of this study is that in contrast to previous studies of metastability, we defined the communities of oscillators directly from the phase-locking data and not from intrinsic connectivity networks. Our so-derived communities are not distinct, specifically mode *ψ*_0_ comprises all other modes. This may be a violation of assumptions for calculating some metrics but we believe that it is more representative of what may actually be happening in the brain, that is, coalitions transiently forming between phase-related communities.

Moving on to communities, previous investigations of Φ*^R^* in fMRI data have used a continuous model to compute the relevant information theoretic variables (Luppi et al., 2020b). In this study we adopted the discrete data model which has been used in computational models of weakly coupled Kuramoto oscillators (Mediano et al., 2022, 2016). We computed Φ*_R_* for an integration timescale from 1-500 TRs and retained the max Φ*^R^* obtained as indicative of integrated information for a specific subject in a specific run. Although there is information in the integration timescale that yielded this Φ*^R^*_*max*_, a maximum statistic test (Novelli et al., 2019) would be required before any inferences may be drawn.

We note that there are a number of differences between our findings and those of (Battaglia et al., 2020). Our stochastic walks were based on instantaneous phase-locking and not on smoothed sliding-window temporal correlation. We used a parcellation with 116 rather than 68 anatomical regions which influences the resulting speeds, and potentially the power-law scaling and fluctuation characteristics. We also did not pool our data as we had sufficient datapoints (1198 TRs) for our calculations. Unlike Battaglia et al. we found that between 40-50% of the HCP subjects exhibited a loss of linearity in power-law scaling in any particular run. In fact, just 7 subjects showed ‘genuine’ power-law scaling over the 4 runs. In a previous study investigating fractal scaling in phase synchronization, fluctuations were averaged over all subjects before determining the scaling component α (Daffertshofer et al., 2018) potentially obscuring loss of linearity in some individual subjects. The lack of linear power-law scaling in individual subjects has been noted before (Botcharova, 2014). We did not investigate the reasons for this lack or loss of linearity although there have been suggestions that this may be due to periodic trends (Hu et al., 2001), non-stationarities (Chen et al., 2002) or non-linear transformations (Chen et al., 2005). Indeed, it has recently been reported that different RSNs exhibit different degrees of non-stationarity (Guan et al., 2020). Unravelling the reasons for loss of linearity is beyond the scope of the present paper, but merits future study.

We did not develop any null models to test the validity of the methodologies employed which may be considered a weakness of this study. However, each of these methodologies has already been validated against null models or with surrogate data (Battaglia et al., 2020; Honari et al., 2020; Mediano et al., 2022). In contrast, there have been few studies that used these methodologies to compare performance across fMRI realizations.

We have just started to explore the relationships between metrics from different conceptualizations of brain functioning. It is clear that there are a number of possible avenues for future research arising from this study. A deeper investigation of power-law linearity differences between subjects and runs for reconfiguration speeds could reveal interesting trait or state correlations. Understanding the relationships between the metrics in general, and with respect to integrated information specifically, poses a challenging task. Unravelling these relationships, potentially with computational models, may provide novel insight into the mechanisms and dynamics of functional connectivity. Finally, applying this battery of metrics to longitudinal or individual life-trajectories could uncover novel relationships that have evaded detection with single methodologies.

## 6 Concluding remarks

Neuromarkers need to demonstrate reliability and interpretability before introduction into a clinical environment. A measure of global metastability, a universal phenomenon across multiple conceptualizations of intrinsic brain activity, was found to be the most representative and stable across multiple fMRI acquisitions of the same subjects. This nonergodicity challenges the use of cross-sectional study designs for dFC. Using concepts and tools from complexity science we have described the metastable behavior of fMRI resting-state activity and our findings are congruent with complex system theory. The inter-relationships between metrics derived from dynamical systems theory, information theory, and information dynamics highlight the simultaneous and balanced tendencies for functional segregation and global integration in the healthy brain. Our battery of metrics may one day help to understand why this balance is lost in psychiatric disorders, or how pharmacological interventions can affect this balance.

## CRediT authorship contribution statement

**Fran Hancock**: Conceptualization; Data curation; Formal analysis; Investigation; Methodology; Software; Visualization; Writing - original draft

**Joana Cabral**: Formal analysis, Methodology; Software; Code Validation; Writing - review & editing.

**Andrea I. Luppi**: Writing - review & editing

**Fernando E. Rosas**: Formal analysis, Writing - review & editing.

**Pedro A.M. Mediano**: Formal analysis, Methodology; Writing - review & editing.

**Ottavia Dipasquale**: Writing - review & editing.

**Federico E. Turkheimer**: Formal analysis; Supervision; Writing - review & editing.

All authors participated in the discussion of the ideas, read and approved the submitted version.

## Abbreviations

fMRI: functional magnetic Resonance Imaging
BOLD: blood oxygen level dependent
FC: Functional Connectivity
dFC: dynamic Functional Connectivity
LEiDA: Leading Eigenvector Dynamic Analysis
DFA: detrended fluctuation analysis

## Acknowledgements

The authors would like to acknowledge the use of the following freely available code:
**MATLAB Toolbox dFCwalk** https://github.com/FunDyn/dFCwalk
**FluctuationAnalysis** https://github.com/marlow17/FluctuationAnalysis
**NEUROMARK** framework http://trendscenter.org/software,
**ICC** Arash Salarian (2021). Intraclass Correlation Coefficient (ICC) (https://www.mathworks.com/matlabcentral/fileexchange/22099-intraclass-correlation-coefficient-icc), MATLAB Central File Exchange. Retrieved August 18, 2021. **corrplot** Wei T, Simko V (2021). R package ’corrplot’: Visualization of a Correlation Matrix. (Version 0.92), https://github.com/taiyun/corrplot.
**lmerTest** (Kuznetsova et al., 2017) https://cran.r-project.org/web/packages/lmerTest/index.html
**performance** (Lüdecke et al., 2021) https://github.com/easystats/performance **permutation_htest_np** https://users.aalto.fi/~eglerean/permutations.html **Shade** Javier Montalt Tordera (2021). Filled area plot (https://www.mathworks.com/matlabcentral/fileexchange/69652-filled-area-plot), MATLAB Central File Exchange. Retrieved November 19, 2021.
**Superbar** Scott Lowe (2021). superbar (https://github.com/scottclowe/superbar), GitHub. Retrieved November 19, 2021.
**Ghost** Attractors https://github.com/jvohryzek/GhostAttractors.

## References

1. Abrol, A., Damaraju, E., Miller, R.L., Stephen, J.M., Claus, E.D., Mayer, A.R., Calhoun, V.D., 2017. Replicability of time-varying connectivity patterns in large resting state fMRI samples. NeuroImage 163, 160–176. https://doi.org/10.1016/j.neuroimage.2017.09.020

2. Alonso Martínez, S., Deco, G., Ter Horst, G.J., Cabral, J., 2020. The Dynamics of Functional Brain Networks Associated With Depressive Symptoms in a Nonclinical Sample. Front. Neural Circuits 14, 570583. https://doi.org/10.3389/fncir.2020.570583

3. Battaglia, D., Boudou, T., Hansen, E.C.A., Lombardo, D., Chettouf, S., Daffertshofer, A., McIntosh, A.R., Zimmermann, J., Ritter, P., Jirsa, V., 2020. Dynamic Functional Connectivity between order and randomness and its evolution across the human adult lifespan. NeuroImage 222, 117156. https://doi.org/10.1016/j.neuroimage.2020.117156

4. Bijsterbosch, J., Harrison, S., Duff, E., Alfaro-Almagro, F., Woolrich, M., Smith, S., 2017. Investigations into within- and between-subject resting-state amplitude variations. Neuroimage 159, 57–69. https://doi.org/10.1016/j.neuroimage.2017.07.014

5. Bijsterbosch, J., Harrison, S.J., Jbabdi, S., Woolrich, M., Beckmann, C., Smith, S., Duff, E.P., 2020. Challenges and future directions for representations of functional brain organization. Nat. Neurosci. 23, 1484–1495. https://doi.org/10.1038/s41593-020-00726-z

6. Botcharova, M., 2014. Modelling and analysis of amplitude, phase and synchrony in human brain activity patterns.

7. Buckner, R.L., Krienen, F.M., Castellanos, A., Diaz, J.C., Yeo, B.T.T., 2011. The organization of the human cerebellum estimated by intrinsic functional connectivity. J. Neurophysiol. 106, 2322–2345. https://doi.org/10.1152/jn.00339.2011

8. Cabral, J., Luckhoo, H., Woolrich, M., Joensson, M., Mohseni, H., Baker, A., Kringelbach, M.L., Deco, G., 2014. Exploring mechanisms of spontaneous functional connectivity in MEG: How delayed network interactions lead to structured amplitude envelopes of band-pass filtered oscillations. NeuroImage 90, 423–435. https://doi.org/10.1016/j.neuroimage.2013.11.047

9. Cabral, J., Vidaurre, D., Marques, P., Magalhães, R., Silva Moreira, P., Miguel Soares, J., Deco, G., Sousa, N., Kringelbach, M.L., 2017. Cognitive performance in healthy older adults relates to spontaneous switching between states of functional connectivity during rest. Sci. Rep. 7, 1–13. https://doi.org/10.1038/s41598-017-05425-7

10. Carhart-Harris, R.L., Leech, R., Hellyer, P.J., Shanahan, M., Feilding, A., Tagliazucchi, E., Chialvo, D.R., Nutt, D., 2014. The entropic brain: a theory of conscious states informed by neuroimaging research with psychedelic drugs. Front. Hum. Neurosci. 8. https://doi.org/10.3389/fnhum.2014.00020

11. Chang, C., Metzger, C.D., Glover, G.H., Duyn, J.H., Heinze, H.-J., Walter, M., 2013. Association between heart rate variability and fluctuations in resting-state functional connectivity. NeuroImage 68, 93–104. https://doi.org/10.1016/j.neuroimage.2012.11.038

12. Chen, Z., Hu, K., Carpena, P., Bernaola-Galvan, P., Stanley, H.E., Ivanov, P.Ch., 2005. Effect of nonlinear filters on detrended fluctuation analysis. Phys. Rev. E 71, 011104. https://doi.org/10.1103/PhysRevE.71.011104

13. Chen, Z., Ivanov, P.Ch., Hu, K., Stanley, H.E., 2002. Effect of nonstationarities on detrended fluctuation analysis. Phys. Rev. E 65, 041107. https://doi.org/10.1103/PhysRevE.65.041107

14. Choe, A.S., Nebel, M.B., Barber, A.D., Cohen, J.R., Xu, Y., Pekar, J.J., Caffo, B., Lindquist, M.A., 2017. Comparing Test-Retest Reliability of Dynamic Functional Connectivity Methods. NeuroImage 158, 155–175. https://doi.org/10.1016/j.neuroimage.2017.07.005

15. Córdova-Palomera, A., Kaufmann, T., Persson, K., Alnæs, D., Doan, N.T., Moberget, T., Lund, M.J., Barca, M.L., Engvig, A., Brækhus, A., Engedal, K., Andreassen, O.A., Selbæk, G., Westlye, L.T., 2017. Disrupted global metastability and static and dynamic brain connectivity across individuals in the Alzheimer’s disease continuum. Sci. Rep. 7, 1–14. https://doi.org/10.1038/srep40268

16. Daffertshofer, A., Ton, R., Kringelbach, M.L., Woolrich, M., Deco, G., 2018. Distinct criticality of phase and amplitude dynamics in the resting brain. NeuroImage, Brain Connectivity Dynamics 180, 442–447. https://doi.org/10.1016/j.neuroimage.2018.03.002

17. Deco, G., Kringelbach, M.L., 2016. Metastability and Coherence: Extending the Communication through Coherence Hypothesis Using A Whole-Brain Computational Perspective. Trends Neurosci. 39, 125–135. https://doi.org/10.1016/j.tins.2016.01.001

18. Deco, G., Kringelbach, M.L., Jirsa, V.K., Ritter, P., 2017. The dynamics of resting fluctuations in the brain: metastability and its dynamical cortical core. Sci. Rep. 7, 3095. https://doi.org/10.1038/s41598-017-03073-5

19. Drew, P.J., Mateo, C., Turner, K.L., Yu, X., Kleinfeld, D., 2020. Ultra-slow Oscillations in fMRI and Resting-State Connectivity: Neuronal and Vascular Contributions and Technical Confounds. Neuron 107, 782–804. https://doi.org/10.1016/j.neuron.2020.07.020

20. Du, Y., Fu, Z., Sui, J., Gao, S., Xing, Y., Lin, D., Salman, M., Abrol, A., Rahaman, M.A., Chen, J., Hong, L.E., Kochunov, P., Osuch, E.A., Calhoun, V.D., 2020. NeuroMark: An automated and adaptive ICA based pipeline to identify reproducible fMRI markers of brain disorders. NeuroImage Clin. 28, 102375. https://doi.org/10.1016/j.nicl.2020.102375

21. Figueroa, C.A., Cabral, J., Mocking, R.J.T., Rapuano, K.M., Hartevelt, T.J. van, Deco, G., Expert, P., Schene, A.H., Kringelbach, M.L., Ruhé, H.G., 2019. Altered ability to access a clinically relevant control network in patients remitted from major depressive disorder. Hum. Brain Mapp. 40, 2771–2786. https://doi.org/10.1002/hbm.24559

22. Gabor, D., 1946. Theory of communication. Proc IEE 93, 429457.

23. Glasser, M.F., Sotiropoulos, S.N., Wilson, J.A., Coalson, T.S., Fischl, B., Andersson, J.L., Xu, J., Jbabdi, S., Webster, M., Polimeni, J.R., Van Essen, D.C., Jenkinson, M., 2013. The minimal preprocessing pipelines for the Human Connectome Project. NeuroImage, Mapping the Connectome 80, 105–124. https://doi.org/10.1016/j.neuroimage.2013.04.127

24. Glerean, E., Salmi, J., Lahnakoski, J.M., Jääskeläinen, I.P., Sams, M., 2012. Functional Magnetic Resonance Imaging Phase Synchronization as a Measure of Dynamic Functional Connectivity. Brain Connect. 2, 91–101. https://doi.org/10.1089/brain.2011.0068

25. Gonzalez-Castillo, J., Kam, J.W., Hoy, C.W., Bandettini, P.A., 2021. How to Interpret Resting-State fMRI: Ask Your Participants. J. Neurosci. 41, 1130–1141.

26. Griffanti, L., Salimi-Khorshidi, G., Beckmann, C.F., Auerbach, E.J., Douaud, G., Sexton, C.E., Zsoldos, E., Ebmeier, K.P., Filippini, N., Mackay, C.E., Moeller, S., Xu, J., Yacoub, E., Baselli, G., Ugurbil, K., Miller, K.L., Smith, S.M., 2014. ICA-based artefact removal and accelerated fMRI acquisition for improved resting state network imaging. NeuroImage 95, 232–247. https://doi.org/10.1016/j.neuroimage.2014.03.034

27. Guan, S., Jiang, R., Bian, H., Yuan, J., Xu, P., Meng, C., Biswal, B., 2020. The Profiles of Non-stationarity and Non-linearity in the Time Series of Resting-State Brain Networks. Front. Neurosci. 14, 493. https://doi.org/10.3389/fnins.2020.00493

28. Haken, H., 1996. Basic Concepts of Synergetics II: Formation of Spatio-temporal Patterns, in: Haken, H. (Ed.), Principles of Brain Functioning: A Synergetic Approach to Brain Activity, Behavior and Cognition, Springer Series in Synergetics. Springer, Berlin, Heidelberg, pp. 149–155. https://doi.org/10.1007/978-3-642-79570-1_11

29. Hellyer, P.J., Scott, G., Shanahan, M., Sharp, D.J., Leech, R., 2015. Cognitive Flexibility through Metastable Neural Dynamics Is Disrupted by Damage to the Structural Connectome. J. Neurosci. 35, 9050–9063. https://doi.org/10.1523/JNEUROSCI.4648-14.2015

30. Hellyer, P.J., Shanahan, M., Scott, G., Wise, R.J.S., Sharp, D.J., Leech, R., 2014. The Control of Global Brain Dynamics: Opposing Actions of Frontoparietal Control and Default Mode Networks on Attention. J. Neurosci. 34, 451–461. https://doi.org/10.1523/JNEUROSCI.1853-13.2014

31. Honari, H., Choe, A.S., Lindquist, M.A., 2021. Evaluating phase synchronization methods in fMRI: A comparison study and new approaches. NeuroImage 228, 117704. https://doi.org/10.1016/j.neuroimage.2020.117704

32. Honari, H., Choe, A.S., Lindquist, M.A., 2020. Evaluating phase synchronization methods in fMRI: a comparison study and new approaches. ArXiv200910126 Cs Eess Stat.

33. Hu, K., Ivanov, P.Ch., Chen, Z., Carpena, P., Eugene Stanley, H., 2001. Effect of trends on detrended fluctuation analysis. Phys. Rev. E 64, 011114. https://doi.org/10.1103/PhysRevE.64.011114

34. Jobst, B.M., Hindriks, R., Laufs, H., Tagliazucchi, E., Hahn, G., Ponce-Alvarez, A., Stevner, A.B.A., Kringelbach, M.L., Deco, G., 2017. Increased Stability and Breakdown of Brain Effective Connectivity During Slow-Wave Sleep: Mechanistic Insights from Whole-Brain Computational Modelling. Sci. Rep. 7, 4634. https://doi.org/10.1038/s41598-017-04522-x

35. Kelso, J.A.S., 1995. Dynamic patterns: The self-organization of brain and behavior, Dynamic patterns: The self-organization of brain and behavior. The MIT Press, Cambridge, MA, US.

36. King, M., Hernandez-Castillo, C.R., Poldrack, R.A., Ivry, R.B., Diedrichsen, J., 2019. Functional boundaries in the human cerebellum revealed by a multi-domain task battery. Nat. Neurosci. 22, 1371–1378. https://doi.org/10.1038/s41593-019-0436-x

37. Koo, T.K., Li, M.Y., 2016. A Guideline of Selecting and Reporting Intraclass Correlation Coefficients for Reliability Research. J. Chiropr. Med. 15, 155–163. https://doi.org/10.1016/j.jcm.2016.02.012

38. Kottaram, A., Johnston, L.A., Cocchi, L., Ganella, E.P., Everall, I., Pantelis, C., Kotagiri, R., Zalesky, A., 2019. Brain network dynamics in schizophrenia: Reduced dynamism of the default mode network. Hum. Brain Mapp. 40, 2212–2228. https://doi.org/10.1002/hbm.24519

39. Kuznetsova, A., Brockhoff, P.B., Christensen, R.H.B., 2017. lmerTest Package: Tests in Linear Mixed Effects Models. J. Stat. Softw. 82, 1–26. https://doi.org/10.18637/jss.v082.i13

40. Landis, J.R., Koch, G.G., 1977. The Measurement of Observer Agreement for Categorical Data. Biometrics 33, 159–174. https://doi.org/10.2307/2529310

41. Laumann, T.O., Snyder, A.Z., Mitra, A., Gordon, E.M., Gratton, C., Adeyemo, B., Gilmore, A.W., Nelson, S.M., Berg, J.J., Greene, D.J., McCarthy, J.E., Tagliazucchi, E., Laufs, H., Schlaggar, B.L., Dosenbach, N.U.F., Petersen, S.E., 2017. On the Stability of BOLD fMRI Correlations. Cereb. Cortex 27, 4719–4732. https://doi.org/10.1093/cercor/bhw265

42. Lord, L.-D., Expert, P., Atasoy, S., Roseman, L., Rapuano, K., Lambiotte, R., Nutt, D.J., Deco, G., Carhart-Harris, R.L., Kringelbach, M.L., Cabral, J., 2019. Dynamical exploration of the repertoire of brain networks at rest is modulated by psilocybin. NeuroImage 199, 127–142. https://doi.org/10.1016/j.neuroimage.2019.05.060

43. Lüdecke, D., Ben-Shachar, M.S., Patil, I., Waggoner, P., Makowski, D., 2021. performance: An R Package for Assessment, Comparison and Testing of Statistical Models. J. Open Source Softw. 6, 3139. https://doi.org/10.21105/joss.03139

44. Luppi, A.I., Mediano, P.A.M., Rosas, F.E., Allanson, J., Pickard, J.D., Carhart-Harris, R.L., Williams, G.B., Craig, M.M., Finoia, P., Owen, A.M., Naci, L., Menon, D.K., Bor, D., Stamatakis, E.A., 2020a. A Synergistic Workspace for Human Consciousness Revealed by Integrated Information Decomposition. bioRxiv 2020.11.25.398081. https://doi.org/10.1101/2020.11.25.398081

45. Luppi, A.I., Mediano, P.A.M., Rosas, F.E., Holland, N., Fryer, T.D., O’Brien, J.T., Rowe, J.B., Menon, D.K., Bor, D., Stamatakis, E.A., 2020b. A synergistic core for human brain evolution and cognition (preprint). Neuroscience. https://doi.org/10.1101/2020.09.22.308981

46. Lurie, D.J., Kessler, D., Bassett, D.S., Betzel, R.F., Breakspear, M., Kheilholz, S., Kucyi, A., Liégeois, R., Lindquist, M.A., McIntosh, A.R., Poldrack, R.A., Shine, J.M., Thompson, W.H., Bielczyk, N.Z., Douw, L., Kraft, D., Miller, R.L., Muthuraman, M., Pasquini, L., Razi, A., Vidaurre, D., Xie, H., Calhoun, V.D., 2020. Questions and controversies in the study of time-varying functional connectivity in resting fMRI. Netw. Neurosci. 4, 30–69. https://doi.org/10.1162/netn_a_00116

47. Mediano, P.A.M., Farah, J.C., Shanahan, M., 2016. Integrated Information and Metastability in Systems of Coupled Oscillators. ArXiv160608313 Q-Bio.

48. Mediano, P.A.M., Rosas, F.E., Farah, J.C., Shanahan, M., Bor, D., Barrett, A.B., 2022. Integrated information as a common signature of dynamical and information-processing complexity. Chaos Interdiscip. J. Nonlinear Sci. 32, 013115. https://doi.org/10.1063/5.0063384

49. Mediano, P.A.M., Rosas, F.E., Luppi, A.I., Carhart-Harris, R.L., Bor, D., Seth, A.K., Barrett, A.B., 2021. Towards an extended taxonomy of information dynamics via Integrated Information Decomposition. ArXiv210913186 Phys. Q-Bio.

50. Newman, M.E.J., 2006. Finding community structure in networks using the eigenvectors of matrices. Phys. Rev. E 74, 036104. https://doi.org/10.1103/PhysRevE.74.036104

51. Noble, S., Scheinost, D., Constable, R.T., 2021. A guide to the measurement and interpretation of fMRI test-retest reliability. Curr. Opin. Behav. Sci. 40, 27–32. https://doi.org/10.1016/j.cobeha.2020.12.012

52. Novelli, L., Wollstadt, P., Mediano, P., Wibral, M., Lizier, J.T., 2019. Large-scale directed network inference with multivariate transfer entropy and hierarchical statistical testing. Netw. Neurosci. Camb. Mass 3, 827–847. https://doi.org/10.1162/netn_a_00092

53. Orban, C., Kong, R., Li, J., Chee, M.W.L., Yeo, B.T.T., 2020. Time of day is associated with paradoxical reductions in global signal fluctuation and functional connectivity. PLOS Biol. 18, e3000602. https://doi.org/10.1371/journal.pbio.3000602

54. Peng C-K, null, Mietus, J., Hausdorff, J.M., Havlin, S., Stanley, H.E., Goldberger, A.L., 1993. Long-range anticorrelations and non-Gaussian behavior of the heartbeat. Phys. Rev. Lett. 70, 1343–1346. https://doi.org/10.1103/PhysRevLett.70.1343

55. Pereda, E., Quiroga, R.Q., Bhattacharya, J., 2005. Nonlinear multivariate analysis of neurophysiological signals. Prog. Neurobiol. 77, 1–37. https://doi.org/10.1016/j.pneurobio.2005.10.003

56. Pierce, J.E., Péron, J., 2020. The basal ganglia and the cerebellum in human emotion. Soc. Cogn. Affect. Neurosci. 15, 599–613. https://doi.org/10.1093/scan/nsaa076

57. Ponce-Alvarez, A., Deco, G., Hagmann, P., Romani, G.L., Mantini, D., Corbetta, M., 2015. Resting-State Temporal Synchronization Networks Emerge from Connectivity Topology and Heterogeneity. PLOS Comput. Biol. 11, e1004100. https://doi.org/10.1371/journal.pcbi.1004100

58. Quian Quiroga, R., Kraskov, A., Kreuz, T., Grassberger, P., 2002. Performance of different synchronization measures in real data: A case study on electroencephalographic signals. Phys. Rev. E 65, 041903. https://doi.org/10.1103/PhysRevE.65.041903

59. Raut, R.V., Snyder, A.Z., Mitra, A., Yellin, D., Fujii, N., Malach, R., Raichle, M.E., 2021. Global waves synchronize the brain’s functional systems with fluctuating arousal. Sci. Adv. 7, eabf2709. https://doi.org/10.1126/sciadv.abf2709

60. Salimi-Khorshidi, G., Douaud, G., Beckmann, C.F., Glasser, M.F., Griffanti, L., Smith, S.M., 2014. Automatic denoising of functional MRI data: Combining independent component analysis and hierarchical fusion of classifiers. NeuroImage 90, 449– 468. https://doi.org/10.1016/j.neuroimage.2013.11.046

61. Schneider, M., Hathway, P., Leuchs, L., Sämann, P.G., Czisch, M., Spoormaker, V.I., 2016. Spontaneous pupil dilations during the resting state are associated with activation of the salience network. NeuroImage 139, 189–201. https://doi.org/10.1016/j.neuroimage.2016.06.011

62. Sendi, M.S.E., Zendehrouh, E., Fu, Z., Liu, J., Du, Y., Mormino, E., Salat, D.H., Calhoun, V.D., Miller, R.L., 2021. Disrupted dynamic functional network connectivity among cognitive control networks in the progression of Alzheimer’s disease. bioRxiv 2020.12.31.424877. https://doi.org/10.1101/2020.12.31.424877

63. Shrout, P.E., Fleiss, J.L., 1979. Intraclass correlations: uses in assessing rater reliability. Psychol. Bull. 86, 420–428. https://doi.org/10.1037//0033-2909.86.2.420

64. Tambini, A., Davachi, L., 2019. Awake reactivation of prior experiences consolidates memories and biases cognition. Trends Cogn. Sci. 23, 876–890.

65. Tognoli, E., Kelso, J.A.S., 2014. The Metastable Brain. Neuron 81, 35–48. https://doi.org/10.1016/j.neuron.2013.12.022

66. Ton, R., Daffertshofer, A., 2016. Model selection for identifying power-law scaling. NeuroImage 136, 215–226. https://doi.org/10.1016/j.neuroimage.2016.01.008

67. Tononi, G., 2004. An information integration theory of consciousness. BMC Neurosci. 5, 42. https://doi.org/10.1186/1471-2202-5-42

68. Turkheimer, F.E., Rosas, F.E., Dipasquale, O., Martins, D., Fagerholm, E.D., Expert, P., Váša, F., Lord, L.-D., Leech, R., 2021. A Complex Systems Perspective on Neuroimaging Studies of Behavior and Its Disorders. The Neuroscientist 1073858421994784. https://doi.org/10.1177/1073858421994784

69. Tzourio-Mazoyer, N., Landeau, B., Papathanassiou, D., Crivello, F., Etard, O., Delcroix, N., Mazoyer, B., Joliot, M., 2002. Automated Anatomical Labeling of Activations in SPM Using a Macroscopic Anatomical Parcellation of the MNI MRI Single- Subject Brain. NeuroImage 15, 273–289. https://doi.org/10.1006/nimg.2001.0978

70. Vaisvilaite, L., Hushagen, V., Grønli, J., Specht, K., 2021. Time-of-Day Effects in Resting-State Functional Magnetic Resonance Imaging: Changes in Effective Connectivity and Blood Oxygenation Level Dependent Signal. Brain Connect. https://doi.org/10.1089/brain.2021.0129

71. Van De Ville, D., Farouj, Y., Preti, M.G., Liégeois, R., Amico, E., 2021. When makes you unique: Temporality of the human brain fingerprint. Sci. Adv. 7, eabj0751. https://doi.org/10.1126/sciadv.abj0751

72. Van Essen, D.C., Smith, S.M., Barch, D.M., Behrens, T.E.J., Yacoub, E., Ugurbil, K., 2013. The WU-Minn Human Connectome Project: An overview. NeuroImage, Mapping the Connectome 80, 62–79. https://doi.org/10.1016/j.neuroimage.2013.05.041

73. Varley, T.F., Luppi, A.I., Pappas, I., Naci, L., Adapa, R., Owen, A.M., Menon, D.K., Stamatakis, E.A., 2020. Consciousness & Brain Functional Complexity in Propofol Anaesthesia. Sci. Rep. 10, 1018. https://doi.org/10.1038/s41598-020-57695-3

74. Vohryzek, J., Deco, G., Cessac, B., Kringelbach, M.L., Cabral, J., 2020. Ghost Attractors in Spontaneous Brain Activity: Recurrent Excursions Into Functionally- Relevant BOLD Phase-Locking States. Front. Syst. Neurosci. 14, 20. https://doi.org/10.3389/fnsys.2020.00020

75. Wildie, M., Shanahan, M., 2012. Metastability and chimera states in modular delay and pulse-coupled oscillator networks. Chaos Interdiscip. J. Nonlinear Sci. 22, 043131. https://doi.org/10.1063/1.4766592

76. Woo, C.-W., Wager, T.D., 2015. Neuroimaging-based biomarker discovery and validation. Pain 156, 1379–1381. https://doi.org/10.1097/j.pain.0000000000000223

77. Xing, X.-X., Zuo, X.-N., 2018. The anatomy of reliability: a must read for future human brain mapping. Sci. Bull. 63, 1606–1607. https://doi.org/10.1016/j.scib.2018.12.010

78. Yang, H., Shew, W.L., Roy, R., Plenz, D., 2012. Maximal Variability of Phase Synchrony in Cortical Networks with Neuronal Avalanches. J. Neurosci. 32, 1061–1072. https://doi.org/10.1523/JNEUROSCI.2771-11.2012

79. Yeo, B.T., Krienen, F.M., Sepulcre, J., Sabuncu, M.R., Lashkari, D., Hollinshead, M., Roffman, J.L., Smoller, J.W., Zöllei, L., Polimeni, J.R., Fischl, B., Liu, H., Buckner, R.L., 2011. The organization of the human cerebral cortex estimated by intrinsic functional connectivity. J. Neurophysiol. 106, 1125–1165. https://doi.org/10.1152/jn.00338.2011

80. Zarghami, T.S., Hossein-Zadeh, G.-A., Bahrami, F., 2020. Deep Temporal Organization of fMRI Phase Synchrony Modes Promotes Large-Scale Disconnection in Schizophrenia. Front. Neurosci. 14, 214. https://doi.org/10.3389/fnins.2020.00214

81. Zhang, J., Kucyi, A., Raya, J., Nielsen, A.N., Nomi, J.S., Damoiseaux, J.S., Greene, D.J., Horovitz, S.G., Uddin, L.Q., Whitfield-Gabrieli, S., 2021. What have we really learned from functional connectivity in clinical populations? NeuroImage 242, 118466. https://doi.org/10.1016/j.neuroimage.2021.118466

82. Zhang, R., Kranz, G.S., Lee, T.M.C., 2019. Functional Connectome from Phase Synchrony at Resting State is a Neural Fingerprint. Brain Connect. 9, 519–528. https://doi.org/10.1089/brain.2018.0657

83. Zhou, Z., Cai, B., Zhang, G., Zhang, A., Calhoun, V.D., Wang, Y.-P., 2020. Prediction and classification of sleep quality based on phase synchronization related whole-brain dynamic connectivity using resting state fMRI. NeuroImage 221, 117190. https://doi.org/10.1016/j.neuroimage.2020.117190

