## Supplementary Material for "Metastability, fractal scaling, and synergistic information processing: what phase relationships reveal about intrinsic brain activity"

### A. Supplementary figures and tables

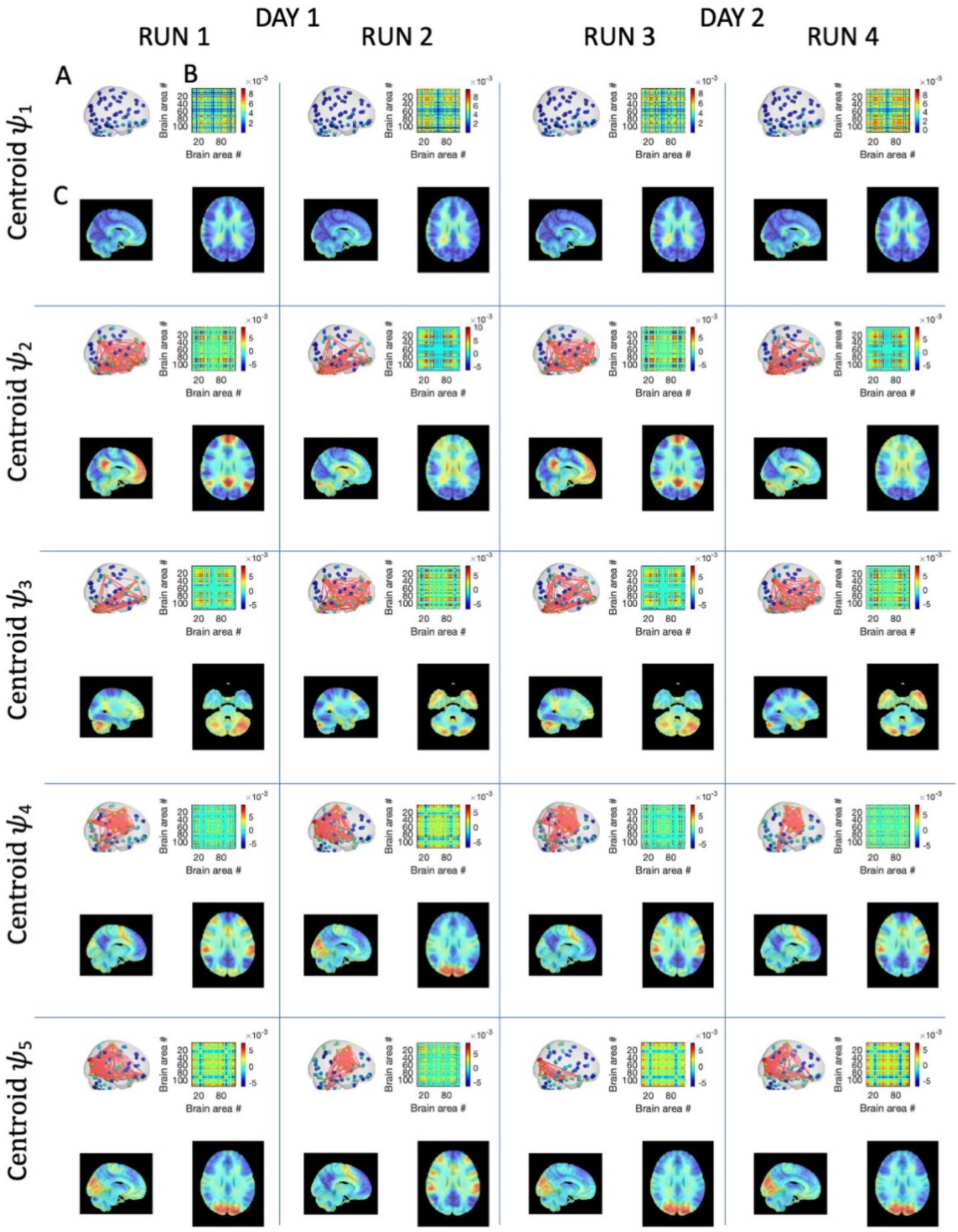

Supplementary Figure 1 Invariant spatiotemporal patterns of phase-locking across 116 regions obtained independently in each of the 4 runs on the same 99 participants.

LEiDA was applied separately to the 4 fMRI runs recorded on 2 consecutive days from 99 participants and the centroids obtained from clustering into K=5 are reported. Each centroid Vc (with size 1xN, with N=116) is represented in three distinct forms: (A) each element Vc(n) is represented as a sphere placed at the center of gravity of the corresponding brain region and its color is scaled according to its value in Vc. Links highlight the network formed by the smallest community of brain areas. (B) the phase-locking matrices computed as the outer product of the centroid vector Vc. (C) Representation of the centroid vector for each mode in 10mm3 voxel space by averaging the eigenvector values over all time instances assigned to a particular cluster/mode. The modes were then plotted over a 1mm3 MNI T1 image.

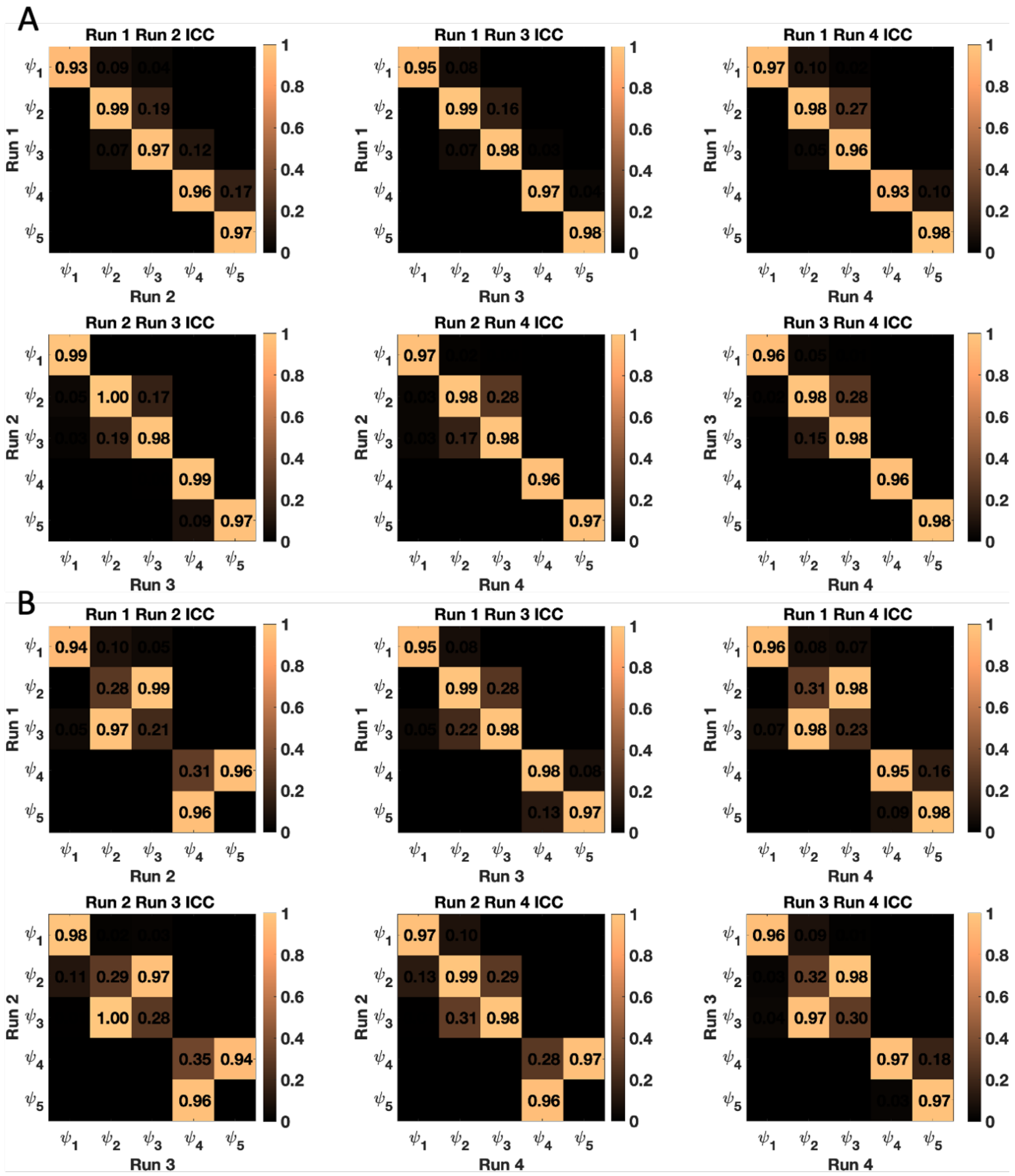

Supplementary Figure 2 Interclass correlation coefficients (ICC) for spatiotemporal patterns of phase-locking extracted with K-means clustering.

(A) Spatiotemporal modes extracted in AAL with 90 regions showed almost perfect agreement between all runs 0.97 > ICC > 0.99. (B) With the inclusion of the cerebellar regions, the order of mode extraction differed between runs. Accounting for this difference run reliability again showed almost perfect agreement 0.94 > ICC > 1.0 in AAL with 116 regions.

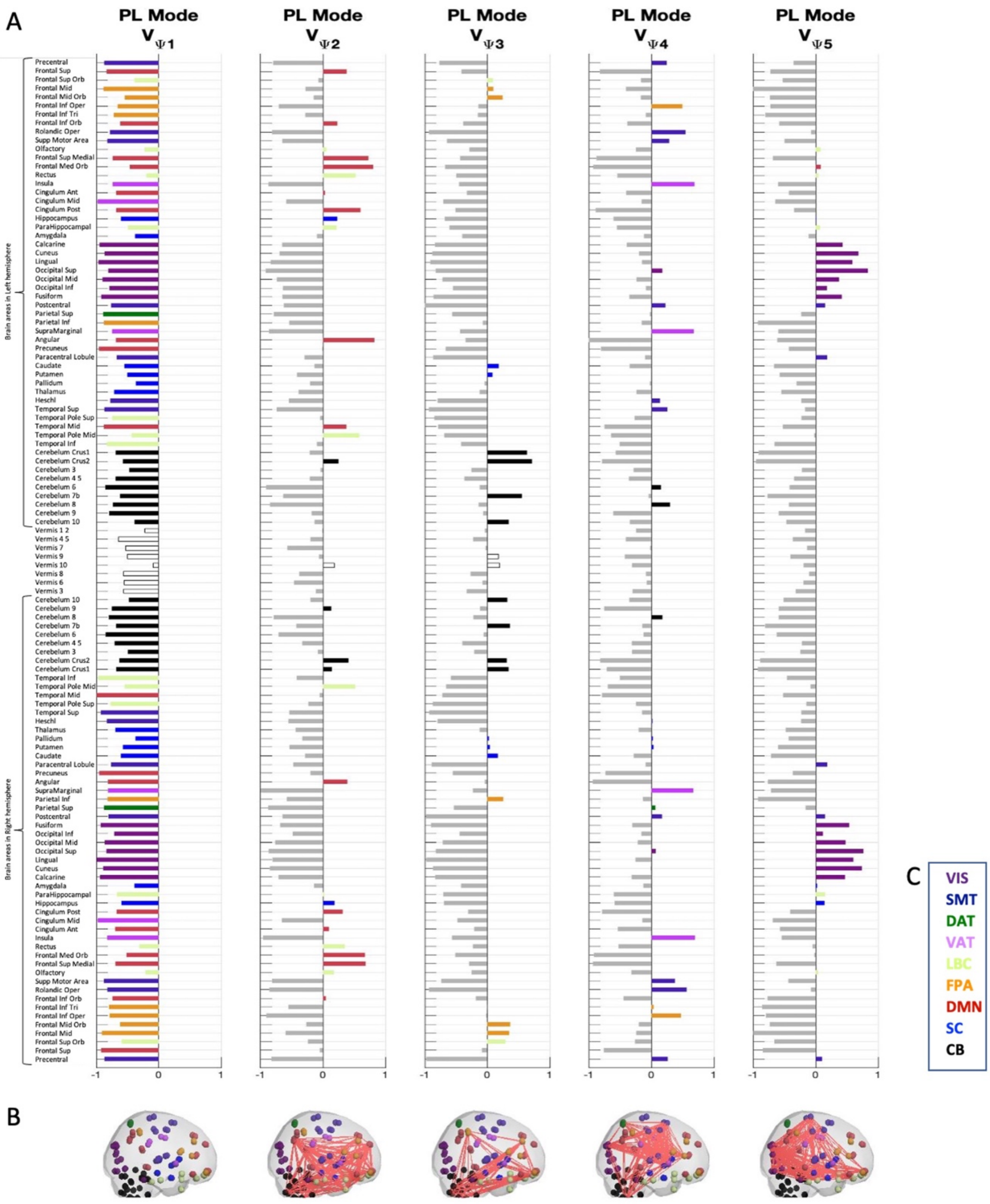

Supplementary Figure 3 Phase-locking modes include sub-elements of Yeo's 7 canonical resting state networks (RSNs).

(A) The bar plot shows the elements in V$_c$ with anti-phase synchronization in mode $\psi_1$, the global mode. (B) The bar plots show the elements in V$_c$ with in-phase synchronization in modes $\psi_2$ to $\psi_5$ color coded to the 7 Yeo RSNs. (B) Representation of the 5 modes obtained when including the cerebellum. We rendered the eigenvectors in cortical space by representing each element as a sphere placed at the center of gravity of the corresponding brain region, and coloring the spheres according to the assigned Yeo resting-state network. We also plot links between the corresponding areas to highlight the network formed by the smallest community of brain areas. (C) Color-codes for the YEO 7 RSNs, visual (VIS), sensory-motor (SM), dorsal attention (DAT), ventral attention (VAT), limbic (LBC) frontal-parietal (FPA), default mode (DMN), the sub-cortical (SC) and cerebella (CB) regions.

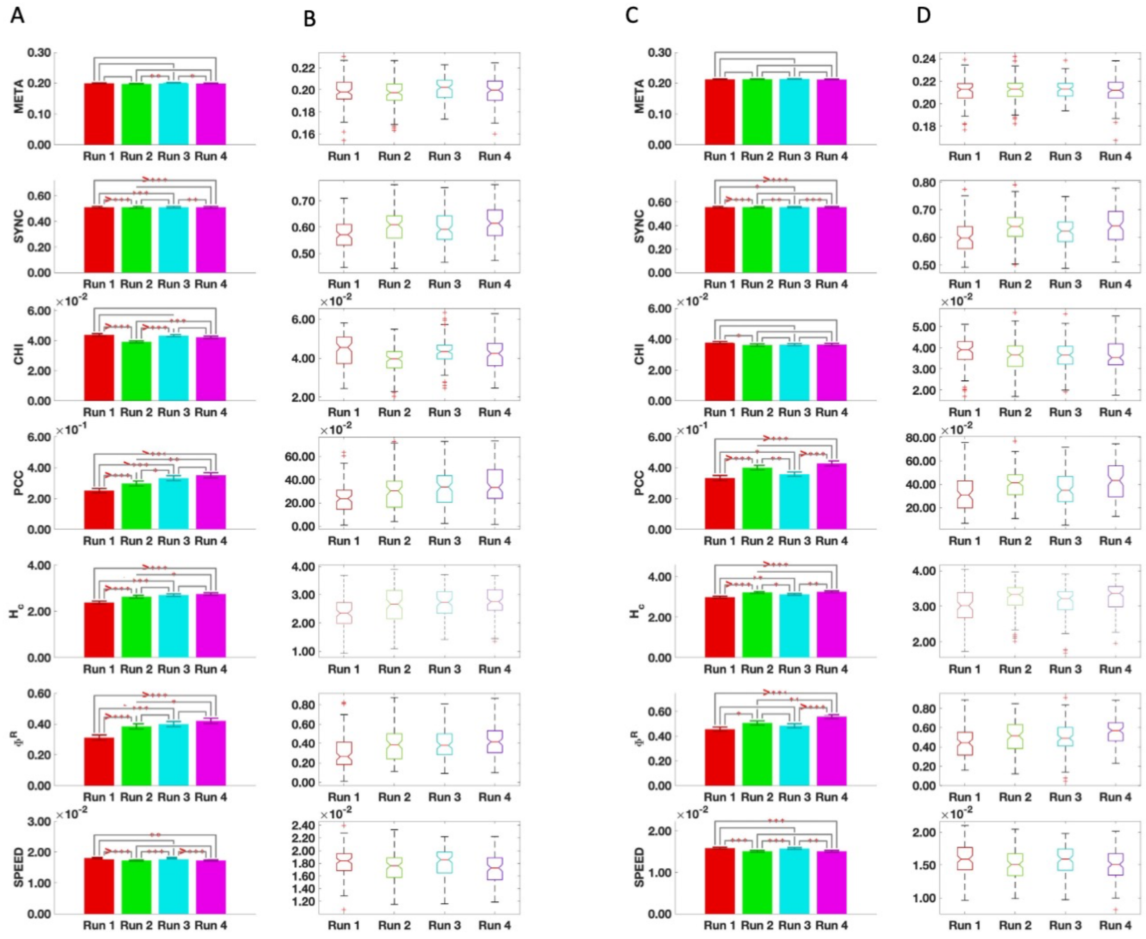

Supplementary Figure 4 Reliability of metastability in parcellations without and with the cerebellar regions

(A) AAL90: The mean values for each metric are shown as bar plots. The *** indicate a statistically significant difference between metrics across the associated runs where * $p < 0.05$, ** $p < 0.01$, *** $p < 0.001$, and >*** $p < 0.0001$. (B) AAL90: The distribution of the global metrics across runs. (C) NEUROMARK: The mean values for each metric are shown as bar plots. The *** indicate a statistically significant difference between individual subject's values of the metric across the associated runs where * $p < 0.05$, ** $p < 0.01$, *** $p < 0.001$, and >*** $p < 0.0001$. (D) NEUROMARK: The distribution of the global metrics across runs. META, metastability, SYNC, synchronization, CHI, chimera index, PCC, phase-coherence coefficient, $H_C$ coalition entropy, $\Phi^R$, integrated information, SPEED, typical reconfiguration speed.

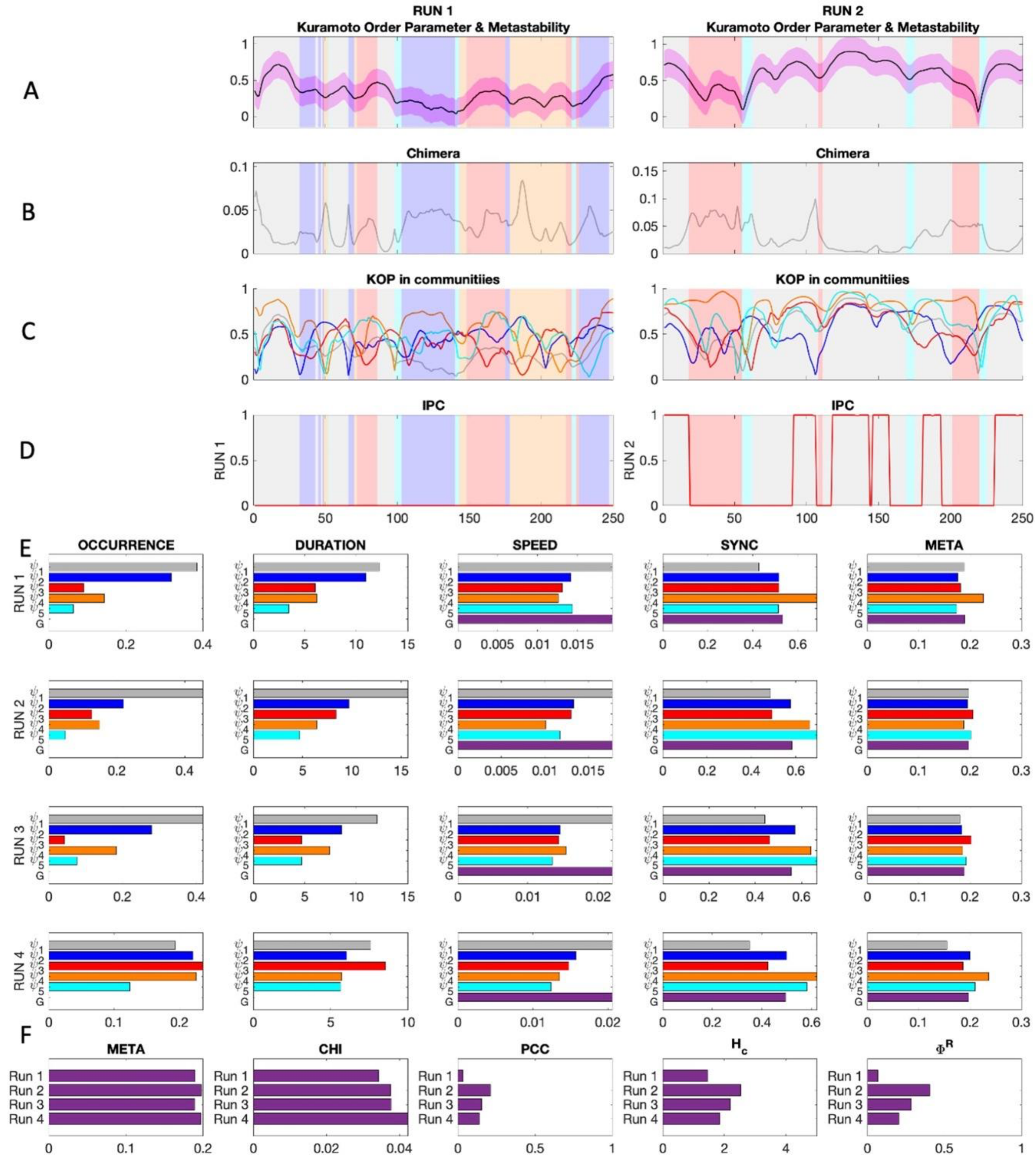

Supplementary Figure 5 Overview of all metrics in all runs for a representative subject. (A) Exemplar snippets from the instantaneous phase synchrony or Kuramoto order parameter time series for each run color-coded to show which mode was dominant at the time. (B) The same as A but for chimeras or cluster synchronization. (C) The evolution of instantaneous synchrony in each of the color-coded modes. (D) The evolution of instantaneous phase-coherence. (E) The values of the mode-specific. (F) The values of the global metrics across all 4 runs. META, metastability, CHI, chimera index, PCC, phase coherence coefficient, $H_c$, coalition entropy, and $\Phi^R$, integrated information

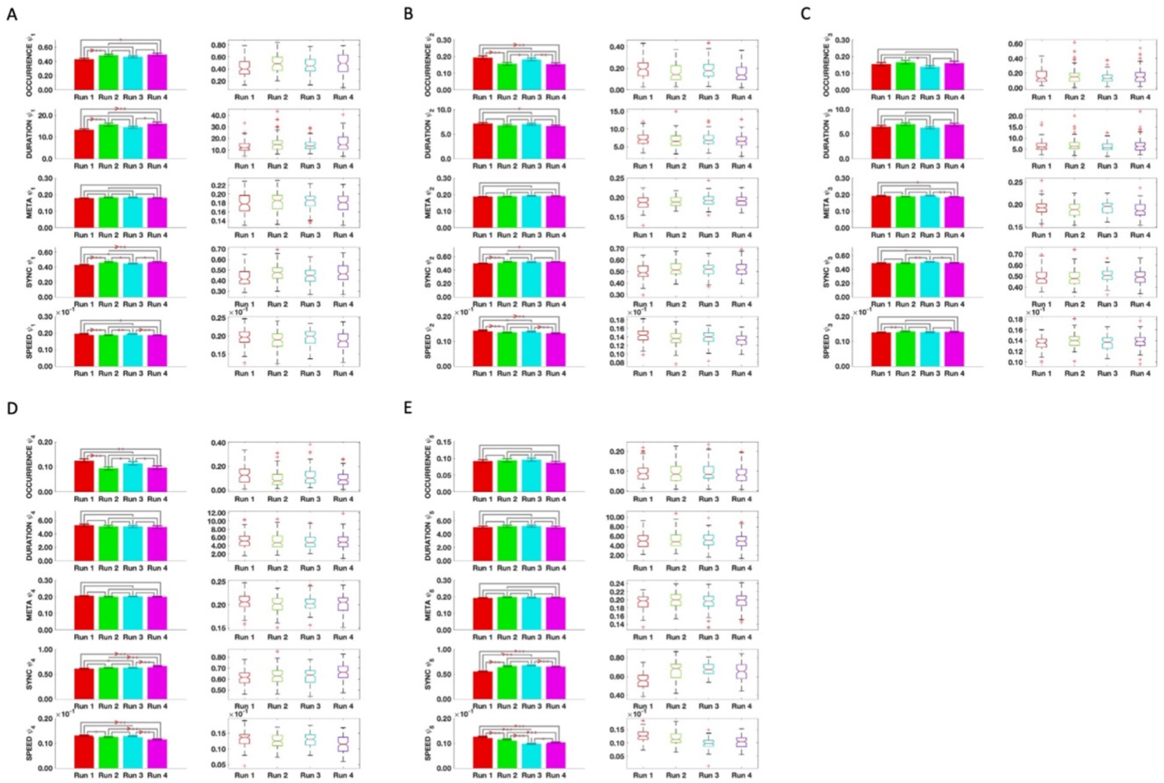

Supplementary Figure 6 Stability of mode-specific metrics in AAL116 for mode $\psi_1 - \psi_5$ across 4 runs. (A-E) Modes $\psi_1 - \psi_5$ respectively. The mean values for each metric are shown as bar plots. The *** indicate a statistically significant difference between the metric across the associated runs where * $p < 0.01$, ** $p < 0.001$, and *** $p < 0.0001$. (B) The distribution of the global metrics across runs. META, metastability, SYNC; synchronization, SPEED, typical speed of reconfiguration.

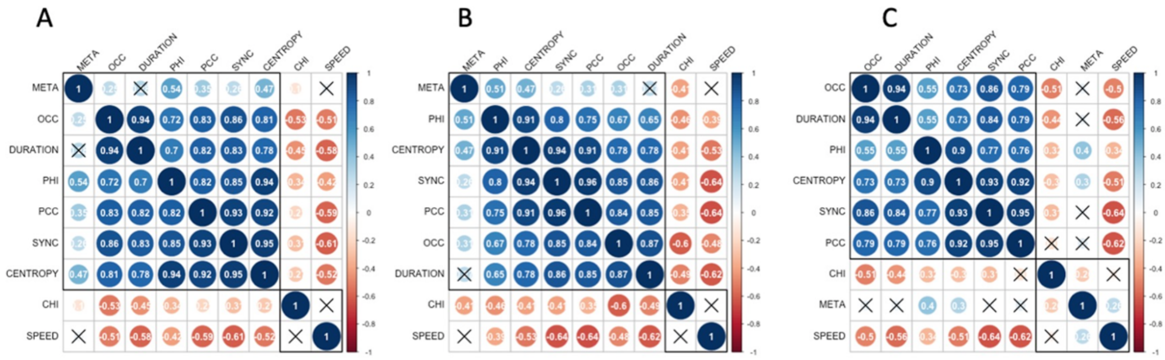

Supplementary Figure 7 Spearman correlation matrices for dFC metrics. (A) Run 2. (B) Run 3. (C) Run 4. META, metastability; OCC, occurrence; PHI, integrated information; PCC, phase coherence coefficient; SYNC, synchronization; CENTROPY, coalition entropy; CHI, chimera index; SPEED, typical reconfiguration speed.

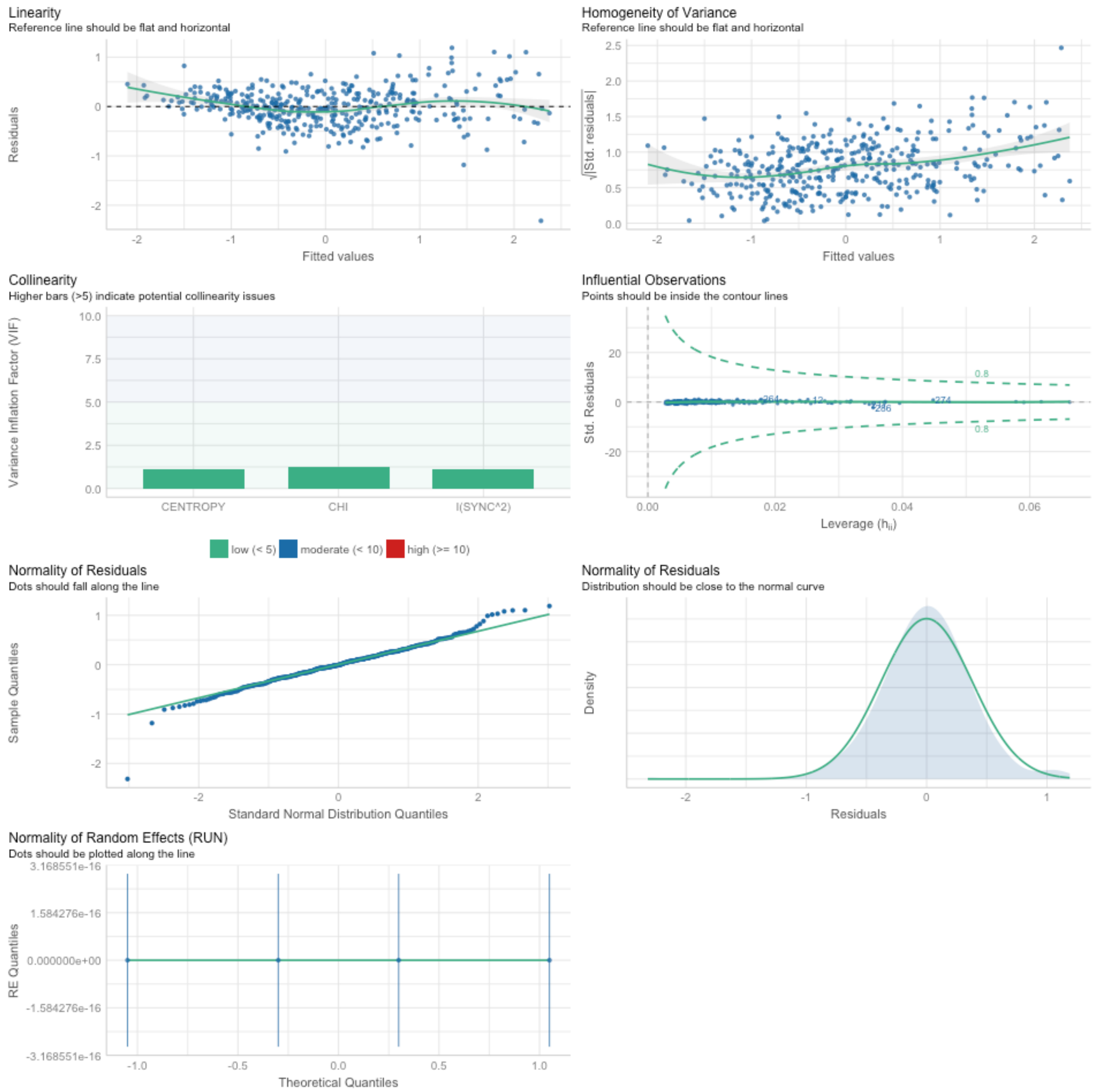

Supplementary Figure 8 Comprehensive visualization of model checks performed on the linear mixed effects regression model to predict Integrated Information from Coalition entropy, Chimera index, and Synchronization. CENTROPY, coalition entropy; CHI, chimera index; SYNC, synchronization.

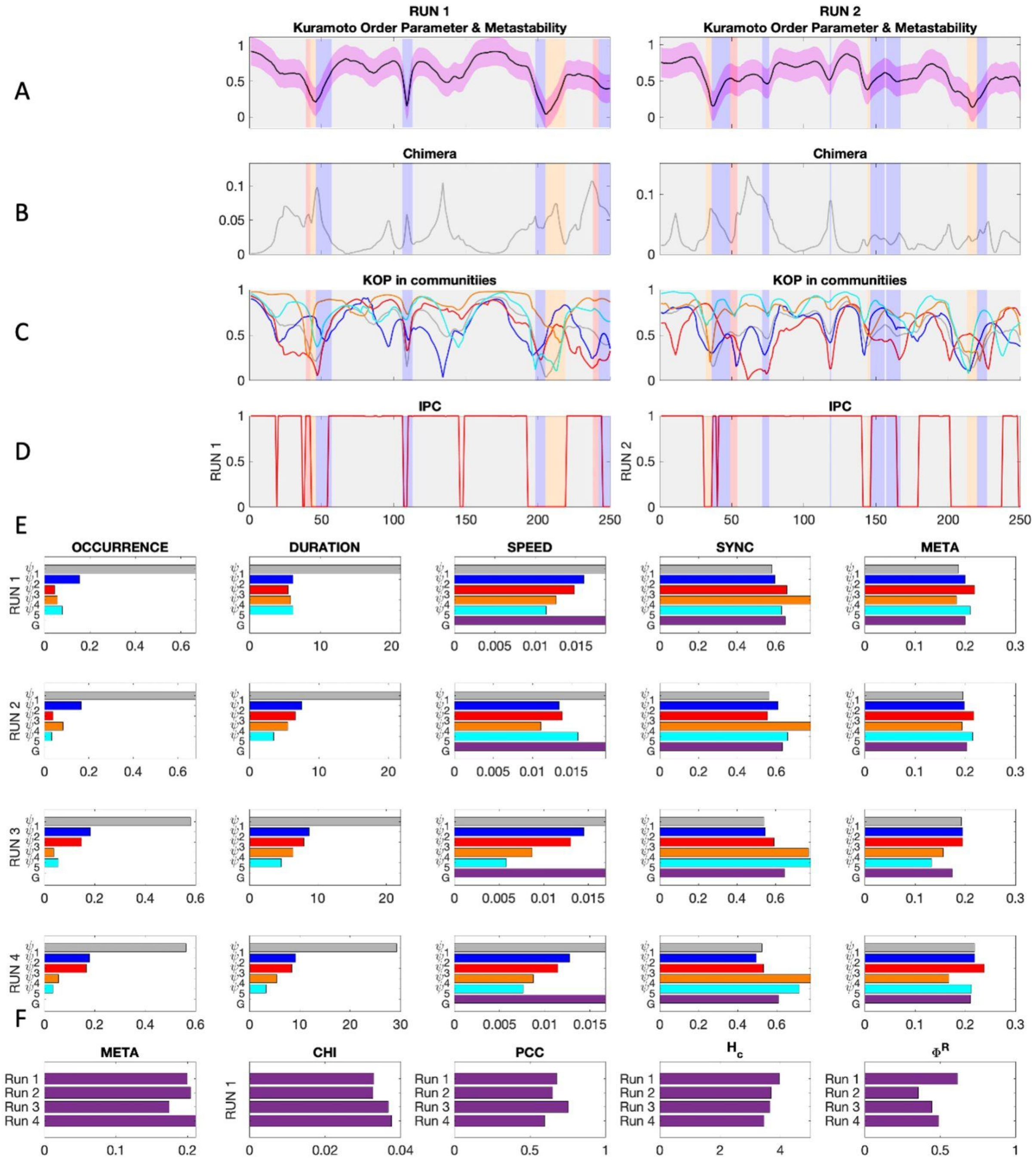

Supplementary Figure 9 Overview of all metrics in all runs for a representative subject with synchronization threshold λ = 0.7.

(A) Exemplar snippets from the instantaneous phase synchrony or Kuramoto order parameter time series for each run color-coded to show which mode was dominant at the time. (B) The same as A but for chimeras or cluster synchronization. (C) The evolution of instantaneous synchrony in each of the color-coded modes. (D) The evolution of instantaneous phase-coherence. (E) The values of the mode-specific. (F) The values of the global metrics across all 4 runs. META, metastability, CHI, chimera index, PCC, phase coherence coefficient, Hc, coalition entropy, and $\Phi^R$, integrated information.

| | | | | | | | | | | | |
|---|---|---|---|---|---|---|---|---|---|---|---|
| 100307.' ':Day 1 | 20:04:46 | rfMRI_REST1_RL | 100307.' ':Day 1 | 20:19:48 | rfMRI_REST1_LR | 100307.' ':Day 2 | 17:34:04 | rfMRI_REST2_RL | 100307.' ':Day 2 | 17:49:00 | rfMRI_REST2_LR |
| 111312.' ':Day 1 | 11:54:45 | rfMRI_REST1_RL | 111312.' ':Day 1 | 12:10:09 | rfMRI_REST1_LR | 111312.' ':Day 2 | 09:09:02 | rfMRI_REST2_RL | 111312.' ':Day 2 | 09:24:11 | rfMRI_REST2_LR |
| 672756.' ':Day 1 | 18:20:30 | rfMRI_REST1_RL | 672756.' ':Day 1 | 18:35:28 | rfMRI_REST1_LR | 672756.' ':Day 2 | 12:32:54 | rfMRI_REST2_RL | 672756.' ':Day 2 | 12:48:01 | rfMRI_REST2_LR |
| 499566.' ':Day 1 | 10:54:28 | rfMRI_REST1_RL | 499566.' ':Day 1 | 11:09:47 | rfMRI_REST1_LR | 499566.' ':Day 2 | 12:06:36 | rfMRI_REST2_RL | 499566.' ':Day 2 | 12:21:48 | rfMRI_REST2_LR |
| 792564.' ':Day 1 | 12:55:18 | rfMRI_REST1_RL | 792564.' ':Day 1 | 13:10:29 | rfMRI_REST1_LR | 792564.' ':Day 2 | 10:48:43 | rfMRI_REST2_RL | 792564.' ':Day 2 | 11:03:44 | rfMRI_REST2_LR |
| 125525.' ':Day 1 | 11:18:29 | rfMRI_REST1_RL | 125525.' ':Day 1 | 11:33:24 | rfMRI_REST1_LR | 125525.' ':Day 2 | 12:49:57 | rfMRI_REST2_RL | 125525.' ':Day 2 | 13:04:50 | rfMRI_REST2_LR |
| 100408.' ':Day 1 | 13:39:44 | rfMRI_REST1_RL | 100408.' ':Day 1 | 13:55:11 | rfMRI_REST1_LR | 100408.' ':Day 2 | 12:20:58 | rfMRI_REST2_LR | 100408.' ':Day 2 | 12:35:55 | rfMRI_REST2_RL |
| 101107.' ':Day 1 | 16:56:38 | rfMRI_REST1_RL | 101107.' ':Day 1 | 17:11:46 | rfMRI_REST1_LR | 101107.' ':Day 2 | 16:05:23 | rfMRI_REST2_LR | 101107.' ':Day 2 | 16:20:35 | rfMRI_REST2_RL |
| 101309.' ':Day 1 | 10:27:56 | rfMRI_REST1_RL | 101309.' ':Day 1 | 10:43:10 | rfMRI_REST1_LR | 101309.' ':Day 2 | 09:47:36 | rfMRI_REST2_LR | 101309.' ':Day 2 | 10:02:56 | rfMRI_REST2_RL |
| 101915.' ':Day 1 | 13:43:08 | rfMRI_REST1_RL | 101915.' ':Day 1 | 13:58:08 | rfMRI_REST1_LR | 101915.' ':Day 2 | 11:32:39 | rfMRI_REST2_LR | 101915.' ':Day 2 | 11:48:06 | rfMRI_REST2_RL |
| 103111.' ':Day 1 | 14:34:20 | rfMRI_REST1_RL | 103111.' ':Day 1 | 14:49:21 | rfMRI_REST1_LR | 103111.' ':Day 2 | 15:31:11 | rfMRI_REST2_LR | 103111.' ':Day 2 | 15:46:06 | rfMRI_REST2_RL |
| 103414.' ':Day 1 | 20:23:35 | rfMRI_REST1_RL | 103414.' ':Day 1 | 20:38:35 | rfMRI_REST1_LR | 103414.' ':Day 3 | 11:59:50 | rfMRI_REST2_LR | 103414.' ':Day 3 | 12:15:25 | rfMRI_REST2_RL |
| 103818.' ':Day 1 | 16:39:22 | rfMRI_REST1_RL | 103818.' ':Day 1 | 16:54:18 | rfMRI_REST1_LR | 103818.' ':Day 2 | 12:38:40 | rfMRI_REST2_LR | 103818.' ':Day 2 | 12:53:53 | rfMRI_REST2_RL |
| 105014.' ':Day 1 | 15:47:19 | rfMRI_REST1_RL | 105014.' ':Day 1 | 16:03:22 | rfMRI_REST1_LR | 105014.' ':Day 2 | 16:25:54 | rfMRI_REST2_LR | 105014.' ':Day 2 | 16:41:07 | rfMRI_REST2_RL |
| 105115.' ':Day 1 | 11:15:50 | rfMRI_REST1_RL | 105115.' ':Day 1 | 11:30:58 | rfMRI_REST1_LR | 105115.' ':Day 2 | 11:11:58 | rfMRI_REST2_LR | 105115.' ':Day 2 | 11:26:58 | rfMRI_REST2_RL |
| 106016.' ':Day 1 | 12:07:17 | rfMRI_REST1_RL | 106016.' ':Day 1 | 12:22:35 | rfMRI_REST1_LR | 106016.' ':Day 2 | 13:26:17 | rfMRI_REST2_LR | 106016.' ':Day 2 | 13:41:19 | rfMRI_REST2_RL |
| 108828.' ':Day 1 | 19:10:58 | rfMRI_REST1_RL | 108828.' ':Day 1 | 19:26:12 | rfMRI_REST1_LR | 108828.' ':Day 2 | 19:33:17 | rfMRI_REST2_LR | 108828.' ':Day 2 | 19:48:54 | rfMRI_REST2_RL |
| 110411.' ':Day 1 | 17:45:43 | rfMRI_REST1_RL | 110411.' ':Day 1 | 18:00:38 | rfMRI_REST1_LR | 110411.' ':Day 2 | 12:09:46 | rfMRI_REST2_LR | 110411.' ':Day 2 | 12:24:40 | rfMRI_REST2_RL |
| 111716.' ':Day 1 | 20:35:15 | rfMRI_REST1_RL | 111716.' ':Day 1 | 20:50:23 | rfMRI_REST1_LR | 111716.' ':Day 2 | 19:53:04 | rfMRI_REST2_LR | 111716.' ':Day 2 | 20:08:00 | rfMRI_REST2_RL |
| 113619.' ':Day 1 | 15:25:06 | rfMRI_REST1_RL | 113619.' ':Day 1 | 15:40:06 | rfMRI_REST1_LR | 113619.' ':Day 2 | 11:02:11 | rfMRI_REST2_LR | 113619.' ':Day 2 | 11:17:05 | rfMRI_REST2_RL |
| 113922.' ':Day 1 | 12:02:21 | rfMRI_REST1_RL | 113922.' ':Day 1 | 12:17:29 | rfMRI_REST1_LR | 113922.' ':Day 2 | 10:56:20 | rfMRI_REST2_LR | 113922.' ':Day 2 | 11:11:31 | rfMRI_REST2_RL |
| 114419.' ':Day 1 | 11:47:44 | rfMRI_REST1_RL | 114419.' ':Day 1 | 12:03:12 | rfMRI_REST1_LR | 114419.' ':Day 2 | 19:06:30 | rfMRI_REST2_LR | 114419.' ':Day 2 | 19:21:27 | rfMRI_REST2_RL |
| 115320.' ':Day 1 | 14:37:24 | rfMRI_REST1_RL | 115320.' ':Day 1 | 14:52:32 | rfMRI_REST1_LR | 115320.' ':Day 2 | 12:39:38 | rfMRI_REST2_LR | 115320.' ':Day 2 | 12:54:43 | rfMRI_REST2_RL |
| 116524.' ':Day 1 | 16:04:16 | rfMRI_REST1_RL | 116524.' ':Day 1 | 16:19:45 | rfMRI_REST1_LR | 116524.' ':Day 2 | 13:12:07 | rfMRI_REST2_LR | 116524.' ':Day 2 | 13:27:04 | rfMRI_REST2_RL |
| 117122.' ':Day 2 | 19:49:45 | rfMRI_REST1_RL | 117122.' ':Day 2 | 20:04:47 | rfMRI_REST1_LR | 117122.' ':Day 3 | 17:46:32 | rfMRI_REST2_LR | 117122.' ':Day 3 | 18:01:25 | rfMRI_REST2_RL |
| 118528.' ':Day 1 | 10:58:28 | rfMRI_REST1_RL | 118528.' ':Day 1 | 11:13:41 | rfMRI_REST1_LR | 118528.' ':Day 2 | 11:57:36 | rfMRI_REST2_LR | 118528.' ':Day 2 | 12:12:52 | rfMRI_REST2_RL |
| 118730.' ':Day 1 | 18:28:50 | rfMRI_REST1_RL | 118730.' ':Day 1 | 18:43:51 | rfMRI_REST1_LR | 118730.' ':Day 2 | 19:03:41 | rfMRI_REST2_LR | 118730.' ':Day 2 | 19:18:37 | rfMRI_REST2_RL |
| 118932.' ':Day 1 | 15:32:42 | rfMRI_REST1_RL | 118932.' ':Day 1 | 15:47:58 | rfMRI_REST1_LR | 118932.' ':Day 2 | 16:19:20 | rfMRI_REST2_LR | 118932.' ':Day 2 | 16:34:18 | rfMRI_REST2_RL |
| 120111.' ':Day 1 | 12:17:40 | rfMRI_REST1_RL | 120111.' ':Day 1 | 12:32:35 | rfMRI_REST1_LR | 120111.' ':Day 2 | 19:21:47 | rfMRI_REST2_LR | 120111.' ':Day 2 | 19:36:58 | rfMRI_REST2_RL |
| 122317.' ':Day 1 | 11:21:36 | rfMRI_REST1_RL | 122317.' ':Day 1 | 11:36:52 | rfMRI_REST1_LR | 122317.' ':Day 2 | 12:44:21 | rfMRI_REST2_LR | 122317.' ':Day 2 | 12:59:42 | rfMRI_REST2_RL |
| 122620.' ':Day 1 | 14:18:18 | rfMRI_REST1_RL | 122620.' ':Day 1 | 14:33:14 | rfMRI_REST1_LR | 122620.' ':Day 2 | 11:01:51 | rfMRI_REST2_LR | 122620.' ':Day 2 | 11:16:49 | rfMRI_REST2_RL |
| 123117.' ':Day 1 | 18:37:12 | rfMRI_REST1_RL | 123117.' ':Day 1 | 18:52:06 | rfMRI_REST1_LR | 123117.' ':Day 2 | 15:48:28 | rfMRI_REST2_LR | 123117.' ':Day 2 | 16:03:21 | rfMRI_REST2_RL |
| 123925.' ':Day 1 | 10:18:51 | rfMRI_REST1_RL | 123925.' ':Day 1 | 10:34:29 | rfMRI_REST1_LR | 123925.' ':Day 2 | 10:38:31 | rfMRI_REST2_LR | 123925.' ':Day 2 | 10:53:37 | rfMRI_REST2_RL |
| 124422.' ':Day 1 | 11:10:22 | rfMRI_REST1_RL | 124422.' ':Day 1 | 11:25:25 | rfMRI_REST1_LR | 124422.' ':Day 2 | 11:21:57 | rfMRI_REST2_LR | 124422.' ':Day 2 | 11:36:56 | rfMRI_REST2_RL |
| 126325.' ':Day 1 | 14:16:26 | rfMRI_REST1_RL | 126325.' ':Day 1 | 14:31:25 | rfMRI_REST1_LR | 126325.' ':Day 2 | 16:07:21 | rfMRI_REST2_LR | 126325.' ':Day 2 | 16:22:14 | rfMRI_REST2_RL |
| 127630.' ':Day 1 | 12:29:22 | rfMRI_REST1_RL | 127630.' ':Day 1 | 12:44:42 | rfMRI_REST1_LR | 127630.' ':Day 2 | 11:55:52 | rfMRI_REST2_LR | 127630.' ':Day 2 | 12:10:57 | rfMRI_REST2_RL |
| 127933.' ':Day 1 | 09:27:56 | rfMRI_REST1_RL | 127933.' ':Day 1 | 09:42:51 | rfMRI_REST1_LR | 127933.' ':Day 2 | 09:37:59 | rfMRI_REST2_LR | 127933.' ':Day 2 | 09:52:54 | rfMRI_REST2_RL |
| 128127.' ':Day 1 | 12:17:44 | rfMRI_REST1_RL | 128127.' ':Day 1 | 12:33:04 | rfMRI_REST1_LR | 128127.' ':Day 2 | 12:14:24 | rfMRI_REST2_LR | 128127.' ':Day 2 | 12:30:03 | rfMRI_REST2_RL |
| 128632.' ':Day 1 | 10:53:12 | rfMRI_REST1_RL | 128632.' ':Day 1 | 11:09:15 | rfMRI_REST1_LR | 128632.' ':Day 2 | 10:51:33 | rfMRI_REST2_LR | 128632.' ':Day 2 | 11:06:46 | rfMRI_REST2_RL |
| 129028.' ':Day 1 | 12:43:26 | rfMRI_REST1_RL | 129028.' ':Day 1 | 12:58:31 | rfMRI_REST1_LR | 129028.' ':Day 2 | 11:51:42 | rfMRI_REST2_LR | 129028.' ':Day 2 | 12:06:51 | rfMRI_REST2_RL |
| 130013.' ':Day 2 | 13:51:00 | rfMRI_REST1_RL | 130013.' ':Day 2 | 14:06:03 | rfMRI_REST1_LR | 130013.' ':Day 3 | 10:05:29 | rfMRI_REST2_LR | 130013.' ':Day 3 | 10:20:26 | rfMRI_REST2_RL |
| 130316.' ':Day 1 | 13:21:33 | rfMRI_REST1_RL | 130316.' ':Day 1 | 13:36:27 | rfMRI_REST1_LR | 130316.' ':Day 2 | 15:21:39 | rfMRI_REST2_LR | 130316.' ':Day 2 | 15:36:35 | rfMRI_REST2_RL |
| 131217.' ':Day 1 | 14:49:53 | rfMRI_REST1_RL | 131217.' ':Day 1 | 15:05:05 | rfMRI_REST1_LR | 131217.' ':Day 2 | 10:04:18 | rfMRI_REST2_LR | 131217.' ':Day 2 | 10:19:18 | rfMRI_REST2_RL |
| 131722.' ':Day 2 | 08:41:53 | rfMRI_REST1_RL | 131722.' ':Day 2 | 08:57:04 | rfMRI_REST1_LR | 131722.' ':Day 3 | 12:28:11 | rfMRI_REST2_LR | 131722.' ':Day 3 | 12:43:41 | rfMRI_REST2_RL |
| 133019.' ':Day 1 | 12:16:53 | rfMRI_REST1_RL | 133019.' ':Day 1 | 12:31:57 | rfMRI_REST1_LR | 133019.' ':Day 2 | 10:59:20 | rfMRI_REST2_LR | 133019.' ':Day 2 | 11:14:42 | rfMRI_REST2_RL |
| 133928.' ':Day 1 | 11:17:51 | rfMRI_REST1_RL | 133928.' ':Day 1 | 11:33:09 | rfMRI_REST1_LR | 133928.' ':Day 2 | 11:49:29 | rfMRI_REST2_LR | 133928.' ':Day 2 | 12:04:44 | rfMRI_REST2_RL |
| 135225.' ':Day 1 | 11:40:36 | rfMRI_REST1_RL | 135225.' ':Day 1 | 11:55:46 | rfMRI_REST1_LR | 135225.' ':Day 2 | 10:59:46 | rfMRI_REST2_LR | 135225.' ':Day 2 | 11:15:55 | rfMRI_REST2_RL |
| 135932.' ':Day 1 | 15:40:11 | rfMRI_REST1_RL | 135932.' ':Day 1 | 15:55:29 | rfMRI_REST1_LR | 135932.' ':Day 2 | 11:19:05 | rfMRI_REST2_LR | 135932.' ':Day 2 | 11:34:13 | rfMRI_REST2_RL |
| 136833.' ':Day 1 | 15:33:04 | rfMRI_REST1_RL | 136833.' ':Day 1 | 15:48:02 | rfMRI_REST1_LR | 136833.' ':Day 2 | 12:29:02 | rfMRI_REST2_LR | 136833.' ':Day 2 | 12:44:15 | rfMRI_REST2_RL |
| 138534.' ':Day 1 | 12:49:36 | rfMRI_REST1_RL | 138534.' ':Day 1 | 13:04:59 | rfMRI_REST1_LR | 138534.' ':Day 2 | 11:33:40 | rfMRI_REST2_LR | 138534.' ':Day 2 | 11:48:34 | rfMRI_REST2_RL |
| 139637.' ':Day 1 | 11:19:04 | rfMRI_REST1_RL | 139637.' ':Day 1 | 11:34:33 | rfMRI_REST1_LR | 139637.' ':Day 2 | 11:18:28 | rfMRI_REST2_LR | 139637.' ':Day 2 | 11:33:58 | rfMRI_REST2_RL |
| 140925.' ':Day 1 | 12:33:47 | rfMRI_REST1_RL | 140925.' ':Day 1 | 12:48:50 | rfMRI_REST1_LR | 140925.' ':Day 2 | 09:58:15 | rfMRI_REST2_LR | 140925.' ':Day 2 | 10:13:14 | rfMRI_REST2_RL |
| 144832.' ':Day 1 | 14:33:29 | rfMRI_REST1_RL | 144832.' ':Day 1 | 14:48:51 | rfMRI_REST1_LR | 144832.' ':Day 2 | 14:09:08 | rfMRI_REST2_LR | 144832.' ':Day 2 | 14:25:14 | rfMRI_REST2_RL |
| 146432.' ':Day 1 | 13:35:59 | rfMRI_REST1_RL | 146432.' ':Day 1 | 13:51:00 | rfMRI_REST1_LR | 146432.' ':Day 2 | 11:28:40 | rfMRI_REST2_LR | 146432.' ':Day 2 | 11:44:02 | rfMRI_REST2_RL |
| 147737.' ':Day 1 | 12:16:17 | rfMRI_REST1_RL | 147737.' ':Day 1 | 12:31:25 | rfMRI_REST1_LR | 147737.' ':Day 2 | 12:12:53 | rfMRI_REST2_LR | 147737.' ':Day 2 | 12:27:50 | rfMRI_REST2_RL |
| 148335.' ':Day 1 | 16:38:15 | rfMRI_REST1_RL | 148335.' ':Day 1 | 16:53:18 | rfMRI_REST1_LR | 148335.' ':Day 2 | 13:57:56 | rfMRI_REST2_LR | 148335.' ':Day 2 | 14:12:54 | rfMRI_REST2_RL |
| 148840.' ':Day 1 | 19:16:09 | rfMRI_REST1_RL | 148840.' ':Day 1 | 19:31:12 | rfMRI_REST1_LR | 148840.' ':Day 2 | 19:08:36 | rfMRI_REST2_LR | 148840.' ':Day 2 | 19:23:31 | rfMRI_REST2_RL |
| 149337.' ':Day 1 | 17:49:53 | rfMRI_REST1_RL | 149337.' ':Day 1 | 18:04:55 | rfMRI_REST1_LR | 149337.' ':Day 2 | 17:04:58 | rfMRI_REST2_LR | 149337.' ':Day 2 | 17:20:14 | rfMRI_REST2_RL |
| 149539.' ':Day 1 | 16:37:38 | rfMRI_REST1_RL | 149539.' ':Day 1 | 16:52:46 | rfMRI_REST1_LR | 149539.' ':Day 2 | 15:08:14 | rfMRI_REST2_LR | 149539.' ':Day 2 | 15:23:24 | rfMRI_REST2_RL |
| 149741.' ':Day 1 | 16:29:01 | rfMRI_REST1_RL | 149741.' ':Day 1 | 16:44:15 | rfMRI_REST1_LR | 149741.' ':Day 2 | 16:11:01 | rfMRI_REST2_LR | 149741.' ':Day 2 | 16:26:00 | rfMRI_REST2_RL |
| 151223.' ':Day 1 | 13:54:03 | rfMRI_REST1_RL | 151223.' ':Day 1 | 14:09:29 | rfMRI_REST1_LR | 151223.' ':Day 2 | 11:17:39 | rfMRI_REST2_LR | 151223.' ':Day 2 | 11:32:47 | rfMRI_REST2_RL |
| 151526.' ':Day 1 | 10:07:31 | rfMRI_REST1_RL | 151526.' ':Day 1 | 10:22:50 | rfMRI_REST1_LR | 151526.' ':Day 2 | 10:28:29 | rfMRI_REST2_LR | 151526.' ':Day 2 | 10:43:48 | rfMRI_REST2_RL |
| 151627.' ':Day 1 | 14:01:20 | rfMRI_REST1_RL | 151627.' ':Day 1 | 14:16:32 | rfMRI_REST1_LR | 151627.' ':Day 2 | 12:09:14 | rfMRI_REST2_LR | 151627.' ':Day 2 | 12:24:24 | rfMRI_REST2_RL |
| 153025.' ':Day 1 | 17:03:35 | rfMRI_REST1_RL | 153025.' ':Day 1 | 17:18:44 | rfMRI_REST1_LR | 153025.' ':Day 2 | 16:53:43 | rfMRI_REST2_LR | 153025.' ':Day 2 | 17:08:59 | rfMRI_REST2_RL |
| 154734.' ':Day 1 | 10:31:22 | rfMRI_REST1_RL | 154734.' ':Day 1 | 10:46:21 | rfMRI_REST1_LR | 154734.' ':Day 2 | 12:09:52 | rfMRI_REST2_LR | 154734.' ':Day 2 | 12:24:48 | rfMRI_REST2_RL |
| 156637.' ':Day 1 | 11:22:19 | rfMRI_REST1_RL | 156637.' ':Day 1 | 11:37:56 | rfMRI_REST1_LR | 156637.' ':Day 2 | 08:37:11 | rfMRI_REST2_LR | 156637.' ':Day 2 | 08:52:21 | rfMRI_REST2_RL |
| 159340.' ':Day 1 | 18:50:51 | rfMRI_REST1_RL | 159340.' ':Day 1 | 19:05:59 | rfMRI_REST1_LR | 159340.' ':Day 2 | 11:25:20 | rfMRI_REST2_LR | 159340.' ':Day 2 | 11:40:51 | rfMRI_REST2_RL |
| 160123.' ':Day 1 | 10:58:00 | rfMRI_REST1_RL | 160123.' ':Day 1 | 11:13:05 | rfMRI_REST1_LR | 160123.' ':Day 2 | 11:11:53 | rfMRI_REST2_LR | 160123.' ':Day 2 | 11:26:57 | rfMRI_REST2_RL |
| 161731.' ':Day 1 | 18:22:10 | rfMRI_REST1_RL | 161731.' ':Day 1 | 18:37:23 | rfMRI_REST1_LR | 161731.' ':Day 2 | 12:44:42 | rfMRI_REST2_LR | 161731.' ':Day 2 | 12:59:48 | rfMRI_REST2_RL |
| 162733.' ':Day 1 | 13:08:35 | rfMRI_REST1_RL | 162733.' ':Day 1 | 13:23:44 | rfMRI_REST1_LR | 162733.' ':Day 2 | 13:02:08 | rfMRI_REST2_LR | 162733.' ':Day 2 | 13:17:04 | rfMRI_REST2_RL |
| 163129.' ':Day 1 | 18:40:27 | rfMRI_REST1_RL | 163129.' ':Day 1 | 18:55:48 | rfMRI_REST1_LR | 163129.' ':Day 2 | 09:41:47 | rfMRI_REST2_LR | 163129.' ':Day 2 | 09:56:50 | rfMRI_REST2_RL |
| 176542.' ':Day 1 | 13:33:06 | rfMRI_REST1_RL | 176542.' ':Day 1 | 13:48:15 | rfMRI_REST1_LR | 176542.' ':Day 2 | 12:07:11 | rfMRI_REST2_LR | 176542.' ':Day 2 | 12:22:16 | rfMRI_REST2_RL |
| 178950.' ':Day 1 | 19:22:04 | rfMRI_REST1_RL | 178950.' ':Day 1 | 19:37:16 | rfMRI_REST1_LR | 178950.' ':Day 2 | 17:33:10 | rfMRI_REST2_LR | 178950.' ':Day 2 | 17:48:12 | rfMRI_REST2_RL |
| 188347.' ':Day 1 | 16:04:32 | rfMRI_REST1_RL | 188347.' ':Day 1 | 16:19:30 | rfMRI_REST1_LR | 188347.' ':Day 2 | 15:24:11 | rfMRI_REST2_LR | 188347.' ':Day 2 | 15:39:06 | rfMRI_REST2_RL |
| 189450.' ':Day 1 | 11:57:38 | rfMRI_REST1_RL | 189450.' ':Day 1 | 12:12:45 | rfMRI_REST1_LR | 189450.' ':Day 2 | 16:25:35 | rfMRI_REST2_LR | 189450.' ':Day 2 | 16:40:35 | rfMRI_REST2_RL |
| 190031.' ':Day 1 | 14:24:01 | rfMRI_REST1_RL | 190031.' ':Day 1 | 14:39:25 | rfMRI_REST1_LR | 190031.' ':Day 2 | 11:09:06 | rfMRI_REST2_LR | 190031.' ':Day 2 | 11:24:24 | rfMRI_REST2_RL |
| 192540.' ':Day 1 | 12:06:24 | rfMRI_REST1_RL | 192540.' ':Day 1 | 12:21:32 | rfMRI_REST1_LR | 192540.' ':Day 2 | 11:19:25 | rfMRI_REST2_LR | 192540.' ':Day 2 | 11:34:25 | rfMRI_REST2_RL |
| 196750.' ':Day 1 | 13:18:12 | rfMRI_REST1_RL | 196750.' ':Day 1 | 13:33:21 | rfMRI_REST1_LR | 196750.' ':Day 2 | 12:54:38 | rfMRI_REST2_LR | 196750.' ':Day 2 | 13:09:39 | rfMRI_REST2_RL |
| 198451.' ':Day 1 | 11:21:30 | rfMRI_REST1_RL | 198451.' ':Day 1 | 11:36:40 | rfMRI_REST1_LR | 198451.' ':Day 2 | 11:04:28 | rfMRI_REST2_LR | 198451.' ':Day 2 | 11:19:31 | rfMRI_REST2_RL |
| 199655.' ':Day 1 | 11:17:36 | rfMRI_REST1_RL | 199655.' ':Day 1 | 11:33:00 | rfMRI_REST1_LR | 199655.' ':Day 2 | 17:52:11 | rfMRI_REST2_LR | 199655.' ':Day 2 | 18:07:18 | rfMRI_REST2_RL |
| 201111.' ':Day 1 | 14:40:34 | rfMRI_REST1_RL | 201111.' ':Day 1 | 14:55:39 | rfMRI_REST1_LR | 201111.' ':Day 2 | 15:34:32 | rfMRI_REST2_LR | 201111.' ':Day 2 | 15:49:34 | rfMRI_REST2_RL |
| 208226.' ':Day 1 | 10:46:34 | rfMRI_REST1_RL | 208226.' ':Day 1 | 11:01:44 | rfMRI_REST1_LR | 208226.' ':Day 2 | 10:57:46 | rfMRI_REST2_LR | 208226.' ':Day 2 | 11:12:49 | rfMRI_REST2_RL |
| 211417.' ':Day 1 | 19:34:52 | rfMRI_REST1_RL | 211417.' ':Day 1 | 19:49:47 | rfMRI_REST1_LR | 211417.' ':Day 2 | 19:05:16 | rfMRI_REST2_LR | 211417.' ':Day 2 | 19:20:11 | rfMRI_REST2_RL |
| 211720.' ':Day 1 | 18:03:44 | rfMRI_REST1_RL | 211720.' ':Day 1 | 18:18:58 | rfMRI_REST1_LR | 211720.' ':Day 2 | 16:04:56 | rfMRI_REST2_LR | 211720.' ':Day 2 | 16:20:06 | rfMRI_REST2_RL |
| 212318.' ':Day 1 | 17:01:21 | rfMRI_REST1_RL | 212318.' ':Day 1 | 17:16:27 | rfMRI_REST1_LR | 212318.' ':Day 3 | 09:02:45 | rfMRI_REST2_LR | 212318.' ':Day 3 | 09:17:43 | rfMRI_REST2_RL |
| 214423.' ':Day 1 | 11:52:19 | rfMRI_REST1_RL | 214423.' ':Day 1 | 12:07:36 | rfMRI_REST1_LR | 214423.' ':Day 2 | 11:02:32 | rfMRI_REST2_LR | 214423.' ':Day 2 | 11:17:36 | rfMRI_REST2_RL |
| 221319.' ':Day 1 | 13:55:37 | rfMRI_REST1_RL | 221319.' ':Day 1 | 14:11:22 | rfMRI_REST1_LR | 221319.' ':Day 2 | 16:25:54 | rfMRI_REST2_LR | 221319.' ':Day 2 | 16:41:04 | rfMRI_REST2_RL |
| 239944.' ':Day 1 | 12:50:43 | rfMRI_REST1_RL | 239944.' ':Day 1 | 13:05:48 | rfMRI_REST1_LR | 239944.' ':Day 2 | 12:05:35 | rfMRI_REST2_LR | 239944.' ':Day 2 | 12:20:41 | rfMRI_REST2_RL |
| 245333.' ':Day 1 | 12:55:04 | rfMRI_REST1_RL | 245333.' ':Day 1 | 13:10:31 | rfMRI_REST1_LR | 245333.' ':Day 2 | 12:23:49 | rfMRI_REST2_LR | 245333.' ':Day 2 | 12:38:54 | rfMRI_REST2_RL |
| 280739.' ':Day 1 | 20:49:14 | rfMRI_REST1_RL | 280739.' ':Day 1 | 21:04:40 | rfMRI_REST1_LR | 280739.' ':Day 2 | 10:01:58 | rfMRI_REST2_LR | 280739.' ':Day 2 | 10:16:52 | rfMRI_REST2_RL |
| 298051.' ':Day 1 | 10:40:41 | rfMRI_REST1_RL | 298051.' ':Day 1 | 10:55:50 | rfMRI_REST1_LR | 298051.' ':Day 2 | 10:31:36 | rfMRI_REST2_LR | 298051.' ':Day 2 | 10:46:35 | rfMRI_REST2_RL |
| 366446.' ':Day 1 | 12:41:06 | rfMRI_REST1_RL | 366446.' ':Day 1 | 12:56:17 | rfMRI_REST1_LR | 366446.' ':Day 2 | 13:40:23 | rfMRI_REST2_LR | 366446.' ':Day 2 | 13:55:39 | rfMRI_REST2_RL |
| 397760.' ':Day 1 | 16:30:06 | rfMRI_REST1_RL | 397760.' ':Day 1 | 16:45:02 | rfMRI_REST1_LR | 397760.' ':Day 2 | 15:37:18 | rfMRI_REST2_LR | 397760.' ':Day 2 | 15:52:14 | rfMRI_REST2_RL |
| 414229.' ':Day 1 | 11:10:13 | rfMRI_REST1_RL | 414229.' ':Day 1 | 11:25:16 | rfMRI_REST1_LR | 414229.' ':Day 2 | 11:06:43 | rfMRI_REST2_LR | 414229.' ':Day 2 | 11:21:48 | rfMRI_REST2_RL |
| 654754.' ':Day 1 | 12:26:39 | rfMRI_REST1_RL | 654754.' ':Day 1 | 12:41:57 | rfMRI_REST1_LR | 654754.' ':Day 2 | 11:00:46 | rfMRI_REST2_LR | 654754.' ':Day 2 | 11:15:55 | rfMRI_REST2_RL |
| 751348.' ':Day 1 | 19:11:05 | rfMRI_REST1_RL | 751348.' ':Day 1 | 19:26:28 | rfMRI_REST1_LR | 751348.' ':Day 2 | 11:26:48 | rfMRI_REST2_LR | 751348.' ':Day 2 | 11:41:57 | rfMRI_REST2_RL |
| 756055.' ':Day 1 | 11:48:57 | rfMRI_REST1_RL | 756055.' ':Day 1 | 12:07:10 | rfMRI_REST1_LR | 756055.' ':Day 2 | 11:00:48 | rfMRI_REST2_LR | 756055.' ':Day 2 | 11:15:50 | rfMRI_REST2_RL |
| 856766.' ':Day 1 | 09:46:18 | rfMRI_REST1_RL | 856766.' ':Day 1 | 10:01:30 | rfMRI_REST1_LR | 856766.' ':Day 2 | 15:43:08 | rfMRI_REST2_LR | 856766.' ':Day 2 | 15:58:08 | rfMRI_REST2_RL |
| 857263.' ':Day 1 | 16:17:25 | rfMRI_REST1_RL | 857263.' ':Day 1 | 16:32:25 | rfMRI_REST1_LR | 857263.' ':Day 2 | 14:53:54 | rfMRI_REST2_LR | 857263.' ':Day 2 | 15:08:58 | rfMRI_REST2_RL |
| 899885.' ':Day 1 | 15:03:14 | rfMRI_REST1_RL | 899885.' ':Day 1 | 15:18:14 | rfMRI_REST1_LR | 899885.' ':Day 2 | 17:06:53 | rfMRI_REST2_LR | 899885.' ':Day 2 | 17:21:57 | rfMRI_REST2_RL |

Supplementary Table S 1 List of HCP U100 subject IDs and respective scan times that were used in this study

### B. Supplementary methods and metrics

#### Dynamical systems metrics

*Fractional occurrence*

For each subject we assigned every timepoint to its closest centroid $V_c$, providing a subject specific mode time-course. This time-course is the basis for the calculation of two statistical metrics as follow:

Fractional occurrence of mode $\psi$ was calculated as

$$\Pi_{\psi}^{(S)} = \frac{1}{T}\sum_{t=1}^{T} \mathcal{X}\left[x^{*}(t) \in R^{\psi}\right] \qquad \text{(S.1)}$$

where $\mathcal{X}$ is a binary event indicator, $T$ is the number of time points in each scan $S$, $x^{*}$ represents the trajectory, or cluster time-series, and $R^{\psi}$ is the defined cluster under consideration.

*Duration*

Duration was calculated as the mean of all consecutive periods spent in a particular mode

$$D_{\psi} = \frac{1}{p_{\psi}}\sum\nolimits_{1}^{p_{\psi}} C_{p_{\psi}} \qquad \text{(S.2)}$$

where $D_{\psi}$ is the duration of mode $\psi$, $p_{\psi}$ is the number of consecutive periods, and $C_{p_{\psi}}$ is the duration of each consecutive period in each scan $S$.

*Stochastic processes - dFC speed analysis*

An unanswered question regarding dFC is whether spatiotemporal patterns change in a discrete or continuous manner over time. K-means clustering yields a distinct mode for each timepoint, but this mode is just the dominant mode, defined to be the mode with the shortest distance to the cluster centroid, and a number of other modes may also contribute to the resulting spatiotemporal pattern at that timepoint. An alternative perspective is to view dFC as a smooth reconfiguration of phase-related connectivity, and to collapse these relations to a point in the space of possible relations. We can then view the evolution of this point as a stochastic exploration of a high-dimensional space. This approach is a direct adaptation of (Battaglia et al., 2020) for phase-locking. The stochastic exploration, or random walk, is quantitatively characterized by its typical reconfiguration speed and path shape (Battaglia et al., 2020).

Following (Battaglia et al., 2020) and (Hansen et al., 2015) we adopted a notion of similarity between $iPL(t)$ matrices based on the Pearson correlation between their upper triangular elements.

$$dPL(t_1, t_2) = corr\big[(UpperTri\big(iPL(t_1)\big), UpperTri(iPL(t_2))\big] \quad \text{(S.3)}$$

The instantaneous global dPL reconfiguration speed at time $t$ is then

$$Speed_{dPL}(t) = 1 - dPL(t_1, t_2) \quad \text{(S.4)}$$

The typical reconfiguration speed is taken as the median of the instantaneous velocity

$$Speed_{typ} = median(Speed_{dPL})_G \quad \text{(S.5)}$$

We used Pearson correlation rather than cosine similarity (Cabral et al., 2017) to retain close alignment with (Battaglia et al., 2020) and (Lombardo et al., 2020). The same calculations were performed for each spatiotemporal mode $\psi$ returned with K-means clustering analogous with (Lombardo et al., 2020).

To visualize the random walk in 3 dimensions, the sequence of time-dependent phase-locking matrices, or *iPL stream*, served as input features into a t-Stochastic Neighborhood Embedding (t-SNE) algorithm (algorithm = exact, distance = Cosine, Perplexity = 50, LearnRate =2000, exaggeration = 4) in MATLAB (MathWorks R2020b) following (Maaten & Hinton, 2008)(Battaglia et al., 2020).

*Detrended fluctuation analysis*

Detrended Fluctuation Analysis (DFA) is a method to detect intrinsic statistical self-affinity embedded in a time series. Self-affinity is a property of fractal timeseries where the similarity is anisotropic. Statistical fractals refer to self-similarity expressed in terms of statistical properties (Eke et al., 2002). DFA infers a self-affinity coefficient through the comparison of detrended mean square fluctuations of the integrated signal over a range of observation scales in a log-log plot. If a genuine power-law exists over a continuous range of scales, then the log-log plot will have an extended linear section, meaning that fluctuations are similar across different temporal scales. So statistically, we have the same fluctuations if we scale the intensity of the signal with the computed DFA coefficient (Eke et al., 2002).

DFA was employed to characterize the shape of the random walk path. Stochastic processes are descriptions of random phenomena evolving over time that are governed by certain laws of probability. Two specific stochastic processes are of relevance for us: the Gaussian process, which represents random noise; and the Wiener process or Brownian motion, which represents a temporally homogenous additive process (Kakihara, 2003). According to the Signal Summation Conversion (SSC) (Eke et al., 2002), the signal (reconfiguration speed in our case) is seen as the realization of either fractional Gaussian noise (fGn) or fractional Brownian motion (fBn). The fBn is non-stationary with independent stationary increments and underlies the computation of DFA. According to SSC, a fGn signal is converted to an fBn through summation before performance of DFA (Eke et al., 2002). We used the fluctuation analysis toolbox of (Ton & Daffertshofer, 2016) available at https://github.com/marlow17/FluctuationAnalysis to perform the DFA.

The time series of instantaneous $dPL$ speed increments was converted into a fBn signal by computing its cumulative sum (Eke et al., 2000):

$$D\left(t_L\right) = \sum\nolimits_{l=1}^{L} Speed_{dPL(t_l)} \qquad \text{(S.6)}$$

Let $K$ denote the number of time series samples split into $Q$ non-overlapping segments $q = 1 \dots Q$ of length $k$, with $Q = K/k$. For each segment $q$, the fluctuation strength $F_q$ was computed as the squared difference between $D\left(t\right)$ and its trend $D^{(trend)}(t)$ (this is the regression line of $D(t)$ over the interval $t = 1 \dots k$ in the linear case).

$$F_q^2(k) = \frac{1}{k}\sum_{l=0}^{k-l}\left[D\left(t_{(q+l)}\right) - D^{(trend)}\left(t_{(q+l)}\right)\right]^2 \qquad \text{(S.7)}$$

This fluctuation strength scales with segment size $k$ in the case of scale-free correlation. On average a linear power law is found in the form of

$$\log F_q\,(k) = \alpha_{DFA} log\ k + C \tag{S.8}$$

The scaling parameter $\alpha_{DFA}$ resembles the Hurst exponent (Metzler et al., 2014) and may be interpreted as follows:

| | |
|---|---|
| $0 < \alpha_{DFA} < 0.5$: $Speed_{dPL}(t)$ | displays anti-persistent fluctuations |
| $\alpha_{DFA} = 0.5$: $Speed_{dPL}(t)$ | displays uncorrelated Gaussian fluctuations |
| $0.5 < \alpha_{DFA} < 1$: $Speed_{dPL}(t)$ | displays persistent fluctuations (approaching pink noise when $\alpha_{DFA}$ is close to 1) |
| $1 \leq \alpha_{DFA}$: $Speed_{dPL}(t)$ | is non-stationary |

*Power-law scaling test*

Returning to the subject of linear power-law scaling, it is imperative to verify that scaling is indeed linear. If this is not the case, the DFA coefficient cannot be interpreted as a scaling exponent (Eke et al., 2000). Following (Ton & Daffertshofer, 2016), we tested the hypothesis of power-law scaling using a Bayesian model comparison approach. This allowed us to fit each subject's DFA log-log plot with 11 different models. Only subjects with an extended linear section over the complete timeseries, returned as a best fit with the linear model, were retained for metric analysis. These subjects were considered to have a 'genuine' $DFA_\alpha$ exponent.

The DFA analysis toolbox evaluates the density of fluctuations over consecutive segments using a kernel source density estimator, and then estimates the log-likelihood for a certain model to generate fluctuations of a given strength (on a log-scale) as a function of log k. Model selection is then performed through computation of the Bayesian Information Criteria (BIC) for each of the tested models defined as

$$BIC = -2log(\mathcal{L}_{\max}) + plog(Q) \quad \text{(S.9)}$$

Where $p$ represents the number of parameters of the model under study, $Q$ the number of intervals with different length $k$, and $\mathcal{L}_{max}$ denotes the maximum value of the likelihood function $\mathcal{L}$ , which quantifies the goodness-of-fit. The model yielding the lowest $BIC$ was selected as the best one, and if it corresponded to model 1 (the linear model), then the returned DFA value was taken as $DFA_{\alpha}$.

In our analysis of 4 runs, we could only confirm linear power-law scaling over the complete timeseries for between 60% to 70% of our 99 subjects in AAL90 parcellation, and 50% to 60% in AAL116. Of these subjects, less than 10 exhibited power-law scaling over all 4 runs. Consequently, we did not include $DFA_{\alpha}$ as a potential reliable metric for dFC analysis, but we did use the average value of 'genuine' $DFA_{\alpha}$ exponents to assess the nature of the fluctuations. We will however, follow up on these intriguing results in a future study, as non-linear power-law scaling, in many cases due to a drop in linearity towards the end of the timeseries, has been found elsewhere in certain subjects (Botcharova, 2014)

### Parameter selection

The toolbox for random walk analysis (Arbabyazd et al., 2020) was developed for dFC computed with temporal correlations and sliding windows. We referred to previous studies using DFA to identify appropriate parameters for use with phase related measures (Botcharova, 2014; Daffertshofer et al., 2018; Eke et al., 2002; Hardstone et al., 2012; Tagliazucchi et al., 2013, 2016; Tetereva et al., 2020; Ton &

Daffertshofer, 2016). Consequently, parameters used with the function FluctuationAnalysis()(Ton & Daffertshofer, 2016) were set as follows: minimum segment length 10 (Tagliazucchi et al., 2013), maximum segment length of Tmax /2 where Tmax is the number of TRs in the scan (Tagliazucchi et al., 2013; Ton & Daffertshofer, 2016), and number of partitions within a segment 20 (Ton & Daffertshofer, 2016).

**Phase synchrony analysis**

Synergetics and Kuramoto

Synergetics is an advanced theory of the phenomena of self-organization where it is proposed that the brain operates close to instabilities otherwise referred to as phase transitions (Haken, 1989). A phase transition occurs when a system undergoes a qualitative change in its macroscopic order. Leveraging the center manifold theorem in dynamical systems theory, the behavior of a system in the vicinity of a phase transition can be approximated by low-dimensional dynamical systems described by a set of order parameter equations. In the Kuramoto model of weakly-coupled interacting limit-cycle oscillators, the collective dynamics of the entire population of oscillators is measured by the macroscopic complex order parameter (Arenas et al., 2008). Here we quantify metastability as the variability in the Kuramoto order parameter (Wildie & Shanahan, 2012) as previous research has demonstrated that this variability is linked to the underlying metastable cluster synchronization in a system of weakly coupled oscillators (Cabral et al., 2011).

Empirical metastability studies to date have used pre-defined regions or seeds from so-called resting-state networks (RSN) to represent communities of oscillators (Hellyer et al., 2014; Lee et al., 2018, 2019; Lord et al., 2019). In contrast, we decided to take a purely data driven approach, using the recurrent modes extracted with K-means clustering to represent communities of oscillators. In addition to the global community, we considered only the regions that were out of phase with the global mode, that is, only the regions $n$ that had positive values in the respective mode centroids $\mathrm{Vc(n)} > 0$. Note that the communities so defined are not distinct but reflect time-varying coalitions among regions.

The Kuramoto order parameters for any mode $\psi$ provide a quantification of the order among the phases over time. Considering the mean of the phases $\theta(r,t)$ across all areas r in a given community $\psi$,

$$Z_{\psi}(t) = \langle e^{i\theta(r,t)} \rangle_{r \in \psi} \tag{S.10}$$

$Z_{\psi}(t)$ is a complex value where its magnitude, $SYNC_{\psi}= |Z_{\psi}(t)|$, provides a quantification of the degree of synchronization of the community at each time $t$, whereas its phase, $PHASE_{\psi}(t) = \arg\left(Z_{\psi}(t)\right)$, indicates the main direction of the phases in the community at each time $t$, which becomes more meaningful as the magnitude of the order parameter increases.

The mean synchrony over time $\langle SYNC_{\psi} \rangle_T$ provides a quantification of the average degree of synchronization of the community over the complete scan duration. $\mathrm{SYNC}_{\psi}$ takes values between 0 and 1, reflecting phase alignment The standard deviation of the synchrony over time, $\mathrm{META}_{\psi} = \sigma( SYNC_{\psi})_T$, provides a measure of metastability of the mode $\psi$ over the complete scan duration.

If we fix time $t$ and estimate the variance of $Z_C(t)$ over the set of all communities C, we obtain an instantaneous measure of how chimera-like the system is at time $t$.

$$CHI_c(t) = var\left(e^{i\theta(c,t)}\right),\ _{c\in C} \tag{S.11}$$

where a measure of the chimera index, an indicator of cluster synchronization, over the complete scan duration is provided by

$$\chi_G = \langle CHI \rangle_{G_T} \tag{S.12}$$

The instantaneous phase-coherence of highly synchronized communities at time $t$ is given by

$$IPC_S = \left|\langle e^{iPHASE_{\psi}(t)} \rangle_{\psi\in S}\right| \tag{S.13}$$

where $\psi \in S$ if $SYNC_{\psi} > \lambda$ at time $t$ , and $|S| > 1$. The synchronization threshold $\lambda$ was set to 0.8 (Wildie & Shanahan, 2012).

Global META was calculated as $\langle \mathrm{META} \rangle_{\psi}$ and global SYNC was calculated as the mean value for the global mode $\psi_1$, $\langle \mathrm{SYNC} \rangle_{\psi_1}$ as this mode includes all regions of all communities. The Phase-Coherence Coefficient ($PCC$) was calculated as the fractional amount of time that $IPC_S$ was non-zero over the complete scan duration.

*Information dynamics analysis*

Metastability has been described as a subtle blend of segregation and integration among brain regions that show tendencies to diverge and function independently, with tendencies to converge and function collectively (Tognoli & Kelso, 2014). The simultaneous presence of these complementary tendencies implies that information shared between regions will either be differentiated to allow for independent processing or integrated for whole brain behavior. Contingent on the premise that the brain can be comprehensively characterized by its information dynamics (Tononi et al., 1994), the brain's current state contains information about its past and future states (in terms of predictability). A measure of integrated information was first proposed (Balduzzi & Tononi, 2008).

$$\Phi = I\left(X_{t;}\, X_{t+1}\right) - \sum_{i=1}^{n} I(X_t^i;\, X_{t+1}^i) \qquad \text{(S.14)}$$

Where $X_t$ denotes the state of the whole system at time $t$; $X_t^i$ denotes the $i^{th}$ part of $X$ at time $t$. $\Phi$ or Integrated Information, captures the information flow between the past and future observed in the whole system $X$, with the flow observed within each of its part, or the 'whole-minus-the-sum' (Mediano et al., 2019). The expression $I\left(X_{t;}\, X_{t+1}\right)$ is the time-delayed mutual information (TDMI) (Barrett & Seth, 2011) and it quantifies the information flow from the past state to the future state. However, this measure is limited in that it only reflects interactions between pairs of variables, that is between a target and a source. To expand on the capabilities for $\Phi$ for multi-source interactions, a framework known as Partial Information Decomposition (PID) was developed by (Williams & Beer, 2010). PID decomposes the total information that two sources give about a target into four distinct '*partial information atoms'* namely *synergistic, redundant*, and 2 *unique* atoms. This framework was further expanded to account for multivariate interactions with $\Phi\mathrm{ID}$ where each $\Phi\mathrm{ID}$ atom is denoted as a pair of PID atoms (Rosas et al., 2019), and subsequently a modified measure $\Phi^R$ was proposed to capture just the synergy and transfer (unique information flow from one component to another) components of integrated information (Mediano et al., 2019, 2021).

The definition of $\Phi$ for a system is the *effective information* $\varphi$ beyond its *Minimum Information Bipartition* (MIB) (Barrett & Seth, 2011). Effective information is a measure of how much better a system $X$ is at predicting its own future after a time $\tau$ when it is considered as whole compared to when it is considered as the sum of two subsystems $M^1$ and $M^2$. Simply put, $\varphi$ indicates how much predictive information is generated by the system over and above the predictive information generated by the two subsystems.

$$\varphi[X;\tau,\mathcal{B}] = I(X_{t-\tau};X_t) - \sum_{k=1}^{2} I(M_{t-\tau}^k;M_t^k) \tag{S.15}$$

where $\mathcal{B} = \{M^1, M^2\}$ is a given bipartition, $I$ is Shannon's mutual information, and $\tau$ is the *integration timescale*.

A system's MIB is the bipartition with lowest $\varphi$, that is, the bipartition with maximal independence and therefore minimal mutual information. And so, we get

$$\Phi[X;\tau] = \varphi[X;\tau,\mathcal{B}^{MIB}] \tag{S.16}$$

$$\mathcal{B}^{MIB} = arg_{\mathcal{B}} min \frac{\varphi[X;\tau,\mathcal{B}]}{K(\mathcal{B})} \tag{S.17}$$

$$K(\mathcal{B}) = min\{H(M^1), H(M^2)\} \tag{S.18}$$

Where $K$ is a normalization factor introduced to avoid biasing $\Phi$ to excessively unbalanced bipartitions (Mediano et al., 2016, 2022).

Returning to $\Phi^R$ , the integrated information without the redundancy components can then be expressed as:

$$\Phi^R[X;\tau] = \varphi[X;\tau,\mathcal{B}^{MIB}] + \min_{i,j} I(M_{t-\tau}^i;M_t^j) \tag{S.19}$$

where the Minimum Mutual Information (MMI) (Barrett, 2015) redundancy function is added back to the formulation.

*Coalition configuration*

As with any information theoretic measure, $\Phi^R$ is substrate-agnostic, that is, the relevant quantity for the calculation of $\Phi^R$ is some informational state of the system, and not the physical state of the system. We therefore define an informational state mapping upon which to calculate $\Phi^R$. To this end, we use the coalition configuration of the system as the informational state reflecting the set of communities that are highly internally synchronized (Mediano et al., 2016).

$$X_t^c = f(x) = \begin{cases} 1, & if \ SYNC_{\psi}(t) > \gamma) \\ 0, & otherwise \end{cases} \quad \text{(S.20)}$$

where $\gamma$ is the *coalition threshold*. The history of the system is now reduced to 5 interdependent binary time series. In this study we set $\gamma$ = 0.8.

*Coalition Entropy*

Although $\langle \mathrm{META} \rangle_{\psi}$ and $\langle CHI \rangle_{G_T}$ detect the occurrence of cluster synchronization, neither indicate if the system visits a small number of chimera configurations repeatedly, or if the system has a large repertoire of such modes. To quantify the diversity of cluster synchronization, we calculate the coalition entropy (Mediano et al., 2016; Shanahan, 2010; Wildie & Shanahan, 2012).

$$H_c = -\sum_{i=1}^{S} p(s_i) log_2 p(s_i) \quad \text{(S.21)}$$

where $S$ is the set of distinct coalitions the system can generate and $p(s)$ is the probability of coalition $s$ arising at any timepoint $t$. With 5 communities $S = 2^5 = 32$.

### C. References

Arbabyazd, L. M., Lombardo, D., Blin, O., Didic, M., Battaglia, D., & Jirsa, V. (2020). Dynamic Functional Connectivity as a complex random walk: Definitions and the dFCwalk toolbox. *MethodsX*, *7*, 101168. https://doi.org/10.1016/j.mex.2020.101168

Arenas, A., Díaz-Guilera, A., Kurths, J., Moreno, Y., & Zhou, C. (2008). Synchronization in complex networks. *Physics Reports*, *469*(3), 93–153. https://doi.org/10.1016/j.physrep.2008.09.002

Balduzzi, D., & Tononi, G. (2008). Integrated Information in Discrete Dynamical Systems: Motivation and Theoretical Framework. *PLOS Computational Biology*, *4*(6), e1000091. https://doi.org/10.1371/journal.pcbi.1000091

Barrett, A. B. (2015). Exploration of synergistic and redundant information sharing in static and dynamical Gaussian systems. *Physical Review E*, *91*(5), 052802. https://doi.org/10.1103/PhysRevE.91.052802

Barrett, A. B., & Seth, A. K. (2011). Practical Measures of Integrated Information for Time-Series Data. *PLOS Computational Biology*, *7*(1), e1001052. https://doi.org/10.1371/journal.pcbi.1001052

Battaglia, D., Boudou, T., Hansen, E. C. A., Lombardo, D., Chettouf, S., Daffertshofer, A., McIntosh, A. R., Zimmermann, J., Ritter, P., & Jirsa, V. (2020). Dynamic Functional Connectivity between order and randomness and its evolution across the human adult lifespan. *NeuroImage*, *222*, 117156. https://doi.org/10.1016/j.neuroimage.2020.117156

Botcharova, M. (2014). *Modelling and analysis of amplitude, phase and synchrony in human brain activity patterns*. https://discovery.ucl.ac.uk/id/eprint/1443453/1/M%20Botcharova%20Thesis%20v9_FINAL.pdf

Cabral, J., Hugues, E., Sporns, O., & Deco, G. (2011). Role of local network oscillations in resting-state functional connectivity. *NeuroImage*, *57*(1), 130–139. https://doi.org/10.1016/j.neuroimage.2011.04.010

Cabral, J., Vidaurre, D., Marques, P., Magalhães, R., Silva Moreira, P., Miguel Soares, J., Deco, G., Sousa, N., & Kringelbach, M. L. (2017). Cognitive performance in healthy older adults relates to spontaneous switching between states of functional connectivity during rest. *Scientific Reports*, *7*(1), 1–13. https://doi.org/10.1038/s41598-017-05425-7

Daffertshofer, A., Ton, R., Kringelbach, M. L., Woolrich, M., & Deco, G. (2018). Distinct criticality of phase and amplitude dynamics in the resting brain. *NeuroImage*, *180*, 442–447. https://doi.org/10.1016/j.neuroimage.2018.03.002

Eke, A., Hermán, P., Bassingthwaighte, J., Raymond, G., Percival, D., Cannon, M., Balla, I., & Ikrényi, C. (2000). Physiological time series: Distinguishing fractal noises from motions. *Pflügers Archiv*, *439*(4), 403–415. https://doi.org/10.1007/s004249900135

Eke, A., Herman, P., Kocsis, L., & Kozak, L. R. (2002). Fractal characterization of complexity in temporal physiological signals. *Physiological Measurement*, *23*(1), R1–R38. https://doi.org/10.1088/0967-3334/23/1/201

Haken, H. (1989). Synergetics: An overview. *Reports on Progress in Physics*, *52*(5), 515–553. https://doi.org/10.1088/0034-4885/52/5/001

Hansen, E. C. A., Battaglia, D., Spiegler, A., Deco, G., & Jirsa, V. K. (2015). Functional connectivity dynamics: Modeling the switching behavior of the resting state. *NeuroImage*, *105*, 525–535. https://doi.org/10.1016/j.neuroimage.2014.11.001

Hardstone, R., Poil, S.-S., Schiavone, G., Jansen, R., Nikulin, V. V., Mansvelder, H. D., & Linkenkaer-Hansen, K. (2012). Detrended Fluctuation Analysis: A Scale-Free View on Neuronal Oscillations. *Frontiers in Physiology*, *0*. https://doi.org/10.3389/fphys.2012.00450

Hellyer, P. J., Shanahan, M., Scott, G., Wise, R. J. S., Sharp, D. J., & Leech, R. (2014). The Control of Global Brain Dynamics: Opposing Actions of Frontoparietal Control and Default Mode Networks on Attention. *Journal of Neuroscience*, *34*(2), 451–461. https://doi.org/10.1523/JNEUROSCI.1853-13.2014

Kakihara, Y. (2003). Stochastic Processes. In R. A. Meyers (Ed.), *Encyclopedia of Physical Science and Technology (Third Edition)* (pp. 105–116). Academic Press. https://doi.org/10.1016/B0-12-227410-5/00739-0

Lee, W. H., Doucet, G. E., Leibu, E., & Frangou, S. (2018). Resting-state network connectivity and metastability predict clinical symptoms in schizophrenia. *Schizophrenia Research*, *201*, 208–216. https://doi.org/10.1016/j.schres.2018.04.029

Lee, W. H., Moser, D. A., Ing, A., Doucet, G. E., & Frangou, S. (2019). Behavioral and Health Correlates of Resting-State Metastability in the Human

Connectome Project. *Brain Topography*, *32*(1), 80–86. https://doi.org/10.1007/s10548-018-0672-5

Lombardo, D., Cassé-Perrot, C., Ranjeva, J.-P., Le Troter, A., Guye, M., Wirsich, J., Payoux, P., Bartrés-Faz, D., Bordet, R., Richardson, J. C., Felician, O., Jirsa, V., Blin, O., Didic, M., & Battaglia, D. (2020). Modular slowing of resting-state dynamic functional connectivity as a marker of cognitive dysfunction induced by sleep deprivation. *NeuroImage*, *222*, 117155. https://doi.org/10.1016/j.neuroimage.2020.117155

Lord, L.-D., Expert, P., Atasoy, S., Roseman, L., Rapuano, K., Lambiotte, R., Nutt, D. J., Deco, G., Carhart-Harris, R. L., Kringelbach, M. L., & Cabral, J. (2019). Dynamical exploration of the repertoire of brain networks at rest is modulated by psilocybin. *NeuroImage*, *199*, 127–142. https://doi.org/10.1016/j.neuroimage.2019.05.060

Maaten, L. V. D., & Hinton, G. E. (2008). Visualizing Data using t-SNE. *Journal of Machine Learning*, *Res. 9 (Nov.)*, 2579–2605.

Mediano, P. A. M., Farah, J. C., & Shanahan, M. (2016). Integrated Information and Metastability in Systems of Coupled Oscillators. *ArXiv:1606.08313 [q-Bio]*. http://arxiv.org/abs/1606.08313

Mediano, P. A. M., Rosas, F., Carhart-Harris, R. L., Seth, A. K., & Barrett, A. B. (2019). Beyond integrated information: A taxonomy of information dynamics phenomena. *ArXiv:1909.02297 [Physics, q-Bio]*. http://arxiv.org/abs/1909.02297

Mediano, P. A. M., Rosas, F. E., Farah, J. C., Shanahan, M., Bor, D., & Barrett, A. B. (2022). Integrated information as a common signature of dynamical and information-processing complexity. *Chaos: An Interdisciplinary Journal of Nonlinear Science*, *32*(1), 013115. https://doi.org/10.1063/5.0063384

Mediano, P. A. M., Rosas, F. E., Luppi, A. I., Carhart-Harris, R. L., Bor, D., Seth, A. K., & Barrett, A. B. (2021). Towards an extended taxonomy of information dynamics via Integrated Information Decomposition. *ArXiv:2109.13186 [Physics, q-Bio]*. http://arxiv.org/abs/2109.13186

Metzler, R., Jeon, J.-H., Cherstvy, A. G., & Barkai, E. (2014). Anomalous diffusion models and their properties: Non-stationarity, non-ergodicity, and ageing at the centenary of single particle tracking. *Phys. Chem. Chem. Phys.*, *16*(44), 24128–24164. https://doi.org/10.1039/C4CP03465A

Rosas, F. E., Mediano, P. A. M., Gastpar, M., & Jensen, H. J. (2019). Quantifying high-order interdependencies via multivariate extensions of the mutual information. *Physical Review E*, *100*(3), 032305. https://doi.org/10.1103/PhysRevE.100.032305

Shanahan, M. (2010). Metastable chimera states in community-structured oscillator networks. *Chaos: An Interdisciplinary Journal of Nonlinear Science*, *20*(1), 013108. https://doi.org/10.1063/1.3305451

Tagliazucchi, E., Chialvo, D. R., Siniatchkin, M., Amico, E., Brichant, J.-F., Bonhomme, V., Noirhomme, Q., Laufs, H., & Laureys, S. (2016). Large-scale signatures of unconsciousness are consistent with a departure from critical dynamics. *Journal of The Royal Society Interface*, *13*(114), 20151027. https://doi.org/10.1098/rsif.2015.1027

Tagliazucchi, E., von Wegner, F., Morzelewski, A., Brodbeck, V., Jahnke, K., & Laufs, H. (2013). Breakdown of long-range temporal dependence in default mode and attention networks during deep sleep. *Proceedings of the National Academy of Sciences*, *110*(38), 15419–15424. https://doi.org/10.1073/pnas.1312848110

Tetereva, A., Kartashov, S., Ivanitsky, A., & Martynova, O. (2020). Variance and Scale-Free Properties of Resting-State Blood Oxygenation Level-Dependent Signal After Fear Memory Acquisition and Extinction. *Frontiers in Human Neuroscience*, *14*, 509075. https://doi.org/10.3389/fnhum.2020.509075

Tognoli, E., & Kelso, J. A. S. (2014). The Metastable Brain. *Neuron*, *81*(1), 35–48. https://doi.org/10.1016/j.neuron.2013.12.022

Ton, R., & Daffertshofer, A. (2016). Model selection for identifying power-law scaling. *NeuroImage*, *136*, 215–226. https://doi.org/10.1016/j.neuroimage.2016.01.008

Tononi, G., Sporns, O., & Edelman, G. M. (1994). A measure for brain complexity: Relating functional segregation and integration in the nervous system. *Proceedings of the National Academy of Sciences*, *91*(11), 5033–5037. https://doi.org/10.1073/pnas.91.11.5033

Wildie, M., & Shanahan, M. (2012). Metastability and chimera states in modular delay and pulse-coupled oscillator networks. *Chaos: An Interdisciplinary Journal of Nonlinear Science*, *22*(4), 043131. https://doi.org/10.1063/1.4766592

Williams, P. L., & Beer, R. D. (2010). Nonnegative Decomposition of Multivariate Information. *ArXiv:1004.2515 [Math-Ph, Physics:Physics, q-Bio]*. http://arxiv.org/abs/1004.2515
